# Precisely-timed dopamine signals establish distinct kinematic representations of skilled movements

**DOI:** 10.1101/2020.07.29.227298

**Authors:** Alexandra Bova, Matt Gaidica, Amy Hurst, Yoshiko Iwai, Daniel K. Leventhal

## Abstract

Brain dopamine is critical for normal motor control, as evidenced by its importance in Parkinson Disease and related disorders. Current hypotheses are that dopamine influences motor control by “invigorating” movements and regulating motor learning. Most evidence for these aspects of dopamine function comes from simple tasks (e.g., lever pressing). Therefore, the influence of dopamine on motor skills requiring multi-joint coordination is unknown. To determine the effects of precisely-timed dopamine manipulations on the performance of a complex, finely coordinated dexterous skill, we optogenetically stimulated or inhibited midbrain dopamine neurons as rats performed a skilled reaching task. We found that reach kinematics and coordination between gross and fine movements progressively changed with repeated manipulations. However, once established, rats transitioned abruptly between aberrant and baseline reach kinematics in a dopamine-dependent manner. These results suggest that precisely-timed dopamine signals have immediate and long-term influences on motor skill performance, distinct from simply “invigorating” movement.

## Introduction

Brain dopamine plays a critical role in motor control. This is most clearly exemplified by the motor symptoms of Parkinson Disease (PD), in which brain dopamine levels are reduced. PD is defined by tremor, rigidity, bradykinesia, and postural instability, which (mostly) respond to dopamine replacement therapy. However, PD patients also experience significant disability from impaired manual dexterity, which causes difficulty with tasks like tying shoelaces, fastening buttons, and handwriting (Pohar & Allyson Jones, 2009). This symptom is distinct from bradykinesia (Foki et al., 2016), but also responds to dopamine replacement (Gebhardt et al., 2008, Lee et al., 2018). Thus, dopamine plays an important, but poorly defined, role in dexterous skill beyond simply regulating movement speed or amplitude.

Two leading hypotheses regarding the role of dopamine in motor control are that it “invigorates” movement and regulates motor learning. The “vigor” hypothesis derives from the exquisite dopa-responsiveness of bradykinesia in PD, and is supported by extensive experimental evidence. Intrastriatal infusion of dopamine agonists increases locomotion, and both electrical and optogenetic stimulation of midbrain dopamine neurons cause contraversive turning (Arbuthnott & Ungerstedt, 1975, Saunders et al., 2018). Dopamine signaling increases near movement onset and acceleration bouts (Coddington & Dudman, 2018, Howe & Dombeck, 2016, Jin & Costa, 2010, Schultz et al., 1983), and is correlated with movement velocity (Barter et al., 2015, Saunders et al., 2018). Conversely, dopamine depletion and dopamine receptor blockade slow movement (Leventhal et al., 2014, Panigrahi et al., 2015). These studies used scalar readouts that reflect “vigor” (e.g., movement velocity or numbers of rotations), and therefore could not assess dopaminergic influences on multi-joint coordination.

Dopaminergic roles in reinforcement learning may contribute to “non-vigor” aspects of motor control. Phasic dopamine release patterns are broadly consistent with “reward prediction error” (RPE) signals, or the difference in value between anticipated and realized behavioral states (Glimcher, 2011). In reinforcement learning models, the RPE is used to adjust subsequent behavior. While the details of dopamine’s role in implicit learning remain to be fully elucidated (Schultz, 2019), dopamine signaling clearly influences synaptic plasticity and alters future behavior (Dowd & Dunnett, 2005, Leventhal et al., 2014, Mohebi et al., 2019, Parker et al., 2016, Shen et al., 2008). Most evidence for “learning” models of dopamine function come from behavioral tasks that require no movement (e.g., classical conditioning, Tobler et al., 2005), simple movements (e.g., lever presses, Parker et al., 2016), or innate movements (e.g., locomotion, Howe & Dombeck, 2016). For the most part, such tasks have discrete outcomes (e.g., push the right or left lever, initiate locomotion or not). However, dopaminergic roles in instrumental and classical conditioning may extend to tasks with more degrees of freedom. In support of this hypothesis, dopamine neuron firing patterns consistent with RPEs (more accurately, performance prediction errors) are observed in songbirds receiving distorted audio feedback (Gadagkar et al., 2016). In mice, rotarod performance worsens gradually during dopamine receptor blockade, and improves gradually when the blockade is released (Beeler et al., 2012). These results could be explained by dopamine reinforcing specific, successful actions (e.g., paw adjustments on the rotarod) to gradually improve performance (Beeler et al., 2013). Nonetheless, the role of dopamine in skilled, dexterous movements requiring precise multi-joint coordination remains unclear.

The goal of this study was to determine the effects of precisely-timed dopaminergic manipulations on a complex, finely coordinated, and relatively unconstrained motor skill. To do this, we optogenetically stimulated or inhibited midbrain dopamine neurons as rats performed a skilled reaching task. In skilled reaching, rats learn the coordinated forelimb and digit movements to reach for, grasp, and consume sugar pellets. Skilled reaching is readily learned by rats over several sessions (Klein et al., 2012, Lemke et al., 2019), requires precise coordination between the forelimb and digits, and is sensitive to dopamine depletion (Hyland et al., 2019, Whishaw et al., 1986). It is therefore an excellent model for assessing dopaminergic contributions to dexterous skill.

By combining skilled reaching, optogenetics, and measurement of 3-dimensional paw/digit kinematics, we addressed the following questions. First, we asked whether dopamine manipulations affect current or subsequent reaches. If dopamine affects only the current movement, reach kinematics should change immediately with dopamine manipulations. Conversely, if dopamine provides a teaching signal for fine motor coordination, reach kinematics should depend on the history of prior dopaminergic activation. Second, we asked how reach kinematics – specifically coordination between forelimb and digit movements – are influenced by dopamine manipulations. If dopamine plays a purely “invigorating” role in movement, altered dopaminergic signaling should affect only the velocity or amplitude of the reaches.

Instead of pure vigor or learning roles for dopamine, we found a complex pattern of dopaminergic influences on skilled reaching. Consistent with a motor learning function, reach kinematics changed gradually with repeated dopamine neuron stimulation or inhibition. In addition to simple kinematic measures (e.g., reach amplitude), coordination between paw advancement and digit movements also changed with repeated stimulation/inhibition. However, once established, rats transitioned between aberrant and baseline reach kinematics within a single trial in a dopamine-dependent manner. These results indicate that dopamine has both immediate and long-term effects on motor control beyond simply invigorating movement, with important implications for understanding dopamine-linked movement disorders.

## Results

We optogenetically stimulated or inhibited substantia nigra pars compacta (SNc) dopamine neurons at specific moments during rat skilled reaching. Tyrosine hydroxylase (TH)-Cre^+^ rats were injected bilaterally with a double-floxed channelrhodopsin (ChR2), archaerhodopsin (Arch), or control EYFP construct into SNc (Figures 1A, 1C, and 1E). Rats were trained on an automated skilled reaching task that allows synchronization of high-speed video with optogenetics (Figure 1D, Bova et al., 2019, Ellens et al., 2016). Following training, optical fibers were implanted over SNc contralateral to the rat’s preferred reaching paw. Immunohistochemistry confirmed that opsin expression was restricted to TH-expressing neurons in SNc projecting to striatum (Figure 1F and Figure 1 – figure supplement 1).

**Figure 1.**
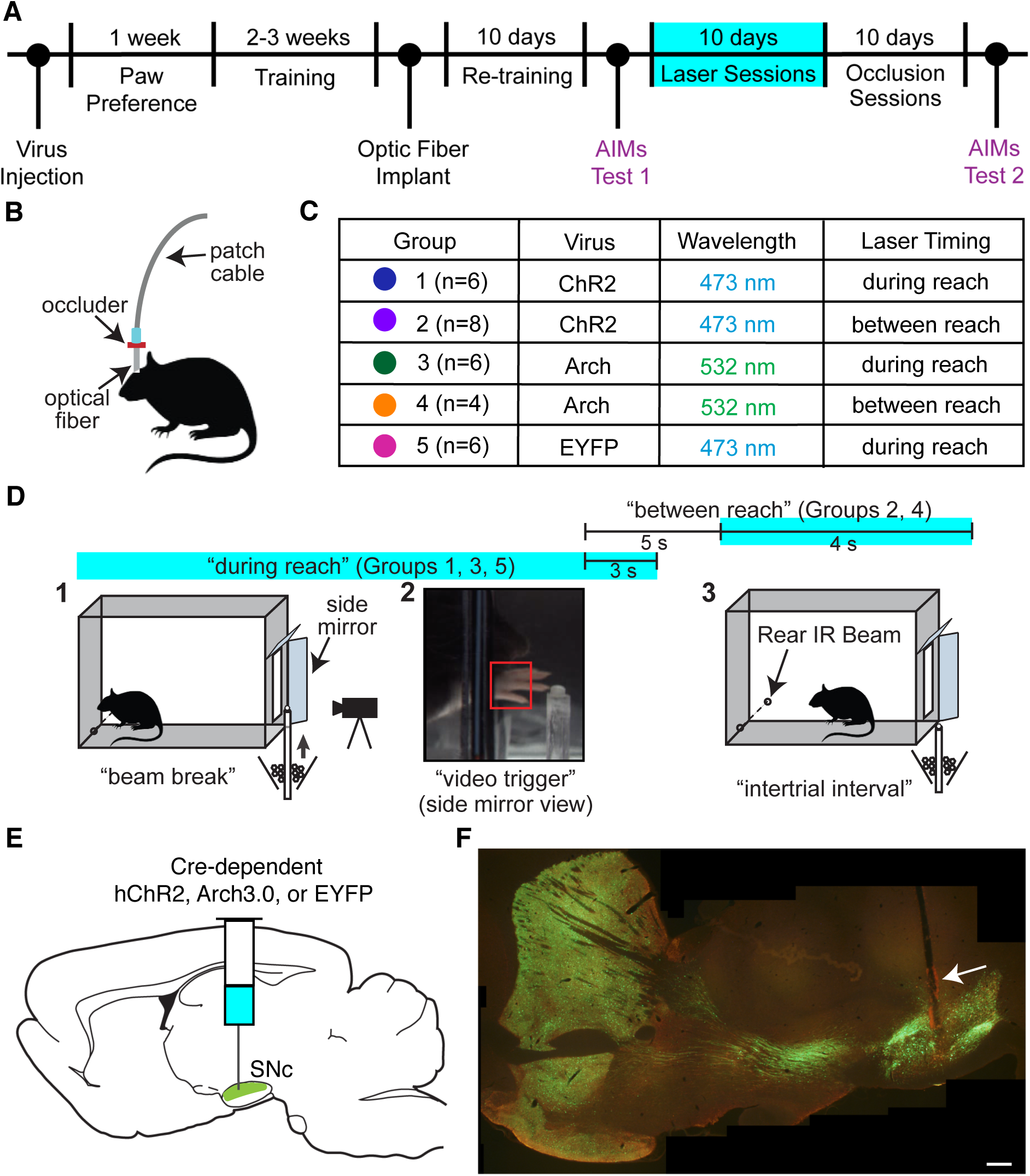
Experimental framework. **(A)** Timeline for a single experiment. AIMs Test – Abnormal Involuntary Movement testing (see “Dopamine neuron stimulation induces context- and history-dependent abnormal involuntary movements”). **(B)** Light was physically occluded from entering the brain by obstructing the connection between the optical fiber and patch cable during “occlusion” sessions. **(C)** Rats were assigned to one of five groups based on virus injected and timing of optogenetic manipulation. *n* is the number of rats included in the analysis for each group (see Materials and Methods). Dot colors correspond with the color used to represent each group in subsequent figures. **(D)** A single skilled reaching trial. 1 – rat breaks IR beam at the back of the chamber to request a sugar pellet (“beam break”). 2 – Real-time analysis detects the paw breaching the reaching slot to trigger 300 fps video from 1 s before to 3.33 s after the trigger event (“video trigger”). 3 – 2 s after the trigger event, the pellet delivery rod resets and the rat can initiate a new trial (“intertrial interval”). Optogenetic manipulations occurred either during reaching (beam break to 3 s after “video trigger”) or between reaches (beginning 5 s after “video trigger” and lasting 4 s). **(E)** Double-floxed ChR2-EYFP, Arch-EYFP, or control EYFP constructs were injected bilaterally into SNc. **(F)** Immunohistochemistry against EYFP showing expression of a fused ChR2-EYFP construct in the nigrostriatal pathway. Optical fibers (arrow) were implanted over SNc contralateral to the rat’s preferred reaching paw. Estimated locations of all fiber tips are shown in Figure 1 – figure supplement 1. Scale bar = 1 mm.

### Altered SNc dopamine neuron activity gradually changes skilled reaching outcomes

We stimulated or inhibited SNc dopamine neurons during every reach for ten 30-minute sessions (Figure 1D, “during reach”). Baseline performance did not differ between groups (Figure 2A, D). Dopamine neuron stimulation did not affect the number of reaches performed (Figure 2B), but caused a progressive decline in reach success (Figure 2E). Reach success rate decreased to about half of baseline performance during the first day of testing, then to nearly 0% for the remainder of “Laser On” sessions. This progressive decline in performance led us to ask whether success rate also changed across trials within individual sessions. Indeed, during the first “Laser On” session, success rate progressively declined (Figure 2G, Figure 2 - figure supplement 1). Furthermore, dopamine-dependent changes in reach success persisted into subsequent sessions. Therefore, reaching performance is dependent on the history of dopamine neuron activation during skilled reaching.

**Figure 2.**
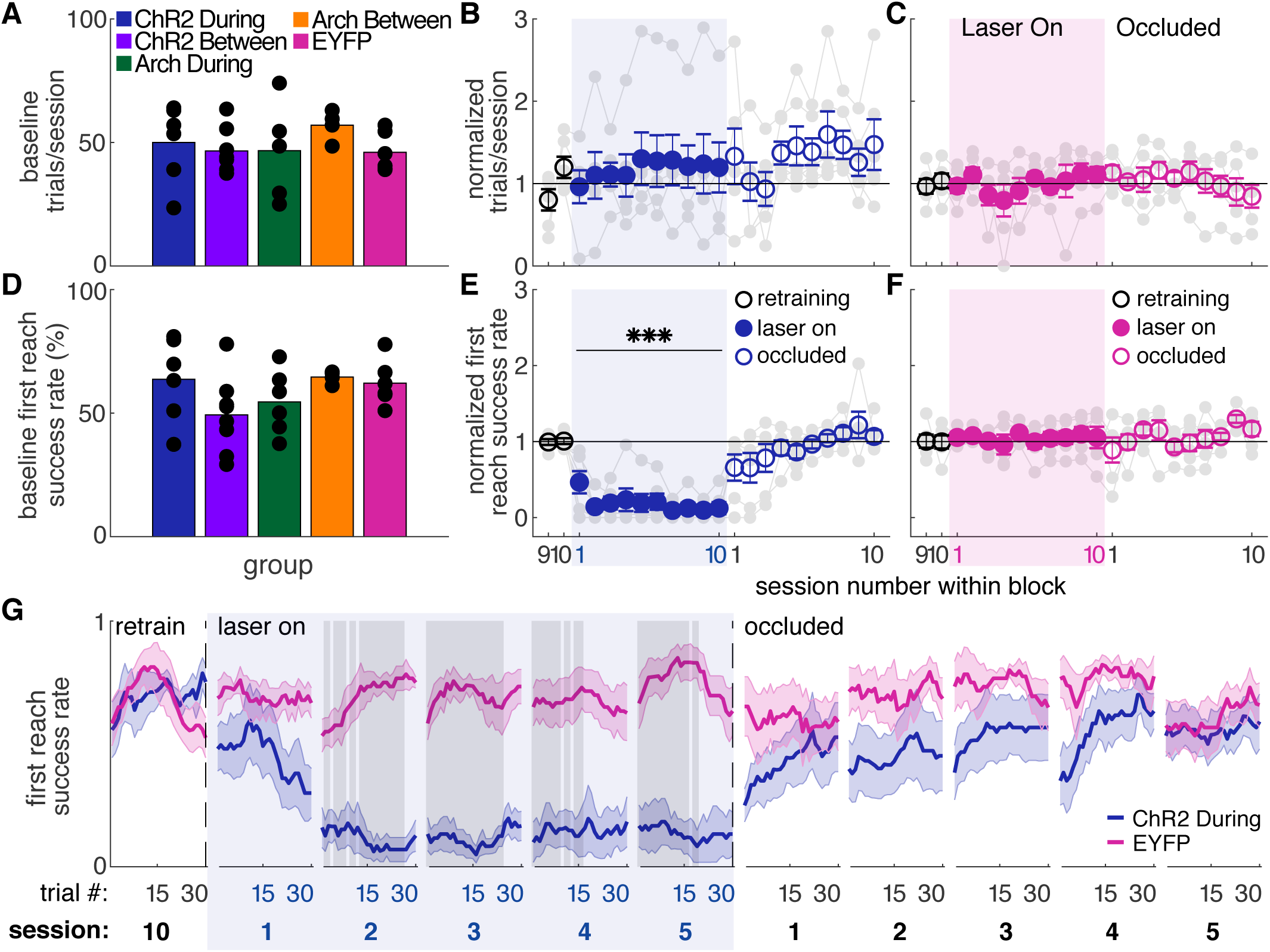
Dopamine neuron stimulation during reaches gradually impairs skilled reaching performance. **(A)** Average number of trials per session over last 2 “retraining” sessions for each group. Black dots represent individual rats. Baseline number of reaches performed did not differ between groups. Kruskal-Wallis Test: χ^2^(4) = 3.94, *P* = 0.41. **(B)** Average number of trials per session divided by the baseline number of trials for “during reach” stimulation. Grey lines represent individual rats. Linear mixed model: effect of laser: *t*(79) = 0.932, *P* = 0.35; interaction between laser and session: *t*(584) = −0.99, *P* = 0.32. **(C)** Same as (B) for control rats injected with an EYFP-only construct. Linear mixed model: effect of laser: *t*(79) = −0.90, *P* = 0.37; interaction between laser and session: *t*(584) = 1.20, *P* = 0.23. **(D)** Average first attempt success rate over the last 2 “retraining” sessions for each group. Black dots represent individual rats. Baseline success rate did not differ between groups. Kruskal-Wallis Test: χ^2^(4) = 6.18, *P* = 0.19. **(E)** Average first attempt success rate divided by baseline success rate for “during reach” stimulation. Linear mixed model: effect of laser: *t*(133) = −3.76, *P* = 2.51×10^-4^; interaction between laser and session: *t*(584) = −1.50, *P* = 0.13. **(F)** Same as (E) for control rats injected with an EYFP-only construct. Linear mixed model: effect of laser: *t*(134) = 0.63, *P* = 0.53; interaction between laser and session: *t*(584) = −0.42, *P* = 0.67. **(G)** Moving average of success rate within individual sessions in the last retraining session, first 5 “laser on” sessions, and first 5 “occlusion” sessions. Shaded grey areas represent statistically significant differences between groups (Wilcoxon rank sum test, *P* < 0.01). Shaded colored areas in (G) and error bars in B-C and E-F represent standard errors of the mean (s.e.m). Data for individual rats are shown in Figure 2 – figure supplement 1. *** indicates p < 0.001 for the laser term in the linear mixed model in panel E.

Because dopamine stimulation during reaching caused a gradual decline in performance, we asked if reaching performance would recover gradually when dopamine stimulation was removed. Animals were tested for an additional 10 days with the same laser stimulation protocol, but with the patch cable-optical fiber junction physically occluded (“occlusion” sessions, Figure 1B). Thus, all cues were identical (e.g., optical shutter noise, visible light) except light penetration into the brain. Reaching performance recovered quickly, but not immediately, to pre-stimulation levels (Figure 2E, G). However, there was significant variability between rats in the rate of recovery (Figure 2 – figure supplement 1). On average, recovery to baseline performance was faster than the decline in performance with initial dopamine stimulation (contrast testing, *t*(583.8) = 2.55, *P* = 0.011). This is further evidence that the history of dopaminergic activation influences subsequent skill execution.

We next asked if dopamine stimulation must occur during reaches to affect success rate. A separate group of ChR2-expressing TH-Cre^+^ rats received laser stimulation during the intertrial interval for a duration matched to “during reach” stimulation (Figure 1D, “between reach”). Dopamine neuron stimulation between reaches had no effect on number of reaches (Figure 3A) or success rate (Figures 3B, C and Figure 3 – figure supplement 1). Therefore, dopamine neuron stimulation must occur as the rat is reaching to affect subsequent reaching performance. This result has two important implications. First, it suggests that skill performance depends on the history of striatal dopamine levels specifically during performance of that skill. Second, it argues against the possibility that the effects of dopamine neuron stimulation are due to the gradual accumulation of striatal dopamine.

**Figure 3.**
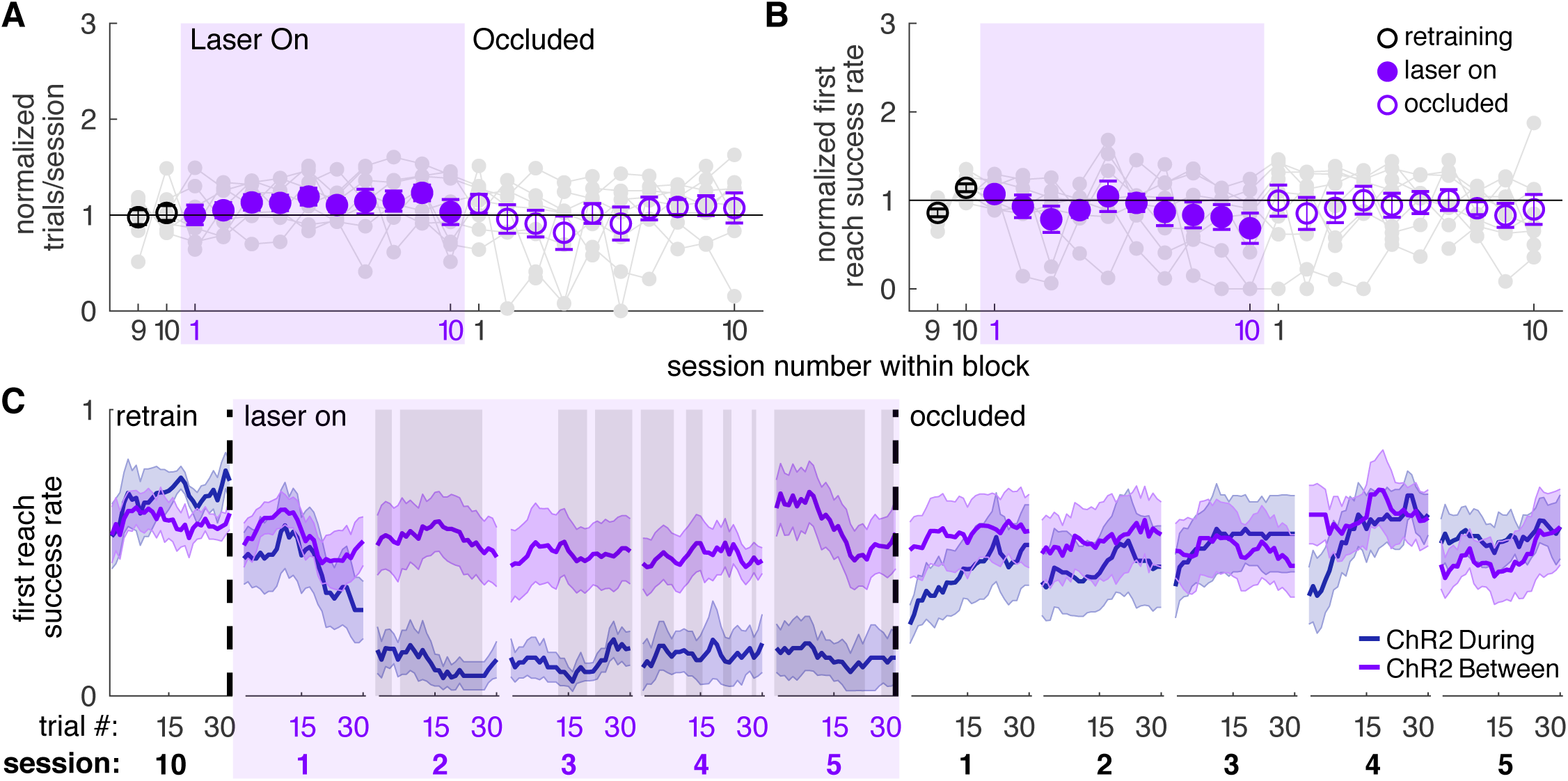
Dopamine neuron stimulation between reaches does not affect skilled reaching performance. **(A)** Average number of trials per session divided by the baseline number of trials for “between reach” stimulation. Grey lines represent individual rats. Linear mixed model: effect of laser: *t*(79) = 1.13, *P* = 0.26; interaction between laser and session: *t*(584) = −0.64, *P* = 0.52. **(B)** Average first attempt success rate divided by baseline success rate for “between reach” stimulation. Linear mixed model: effect of laser: *t*(133) = −0.29, *P* = 0.78; interaction between laser and session: *t*(584) = −0.94, *P* = 0.35. **(C)** Moving average of success rate within individual sessions in the last retraining session, first 5 “laser on” sessions, and first 5 “occlusion” sessions. “During reach” data from Figure 2 are shown for comparison. Shaded grey areas represent trials with a statistically significant difference between groups (Wilcoxon rank sum text, *P* < 0.01). Data for individual rats are shown in Figure 3 – figure supplement 1. Shaded colored areas in C and error bars in A-B represent s.e.m.

Dopamine neuron inhibition during reaching did not affect success rate (Figures 4C, E and Figure 4 – figure supplement 2). However, dopamine neuron inhibition significantly decreased the number of reaches per session (Figure 4A), consistent with a role for midbrain dopamine in motivation to work for rewards (Palmiter, 2008, Salamone & Correa, 2012). This effect was also gradual, with rats progressively performing fewer reaches across sessions. Dopamine neuron inhibition between reaches had no effect on success rate (Figure 4D and Figure 4 – figure supplement 1) or the number of reaches performed in each session (Figure 4B). Control rats injected with constructs expressing EYFP but no opsin did not experience any changes in task performance (Figures 2C, F, G and Figure 2 – figure supplement 1).

**Figure 4.**
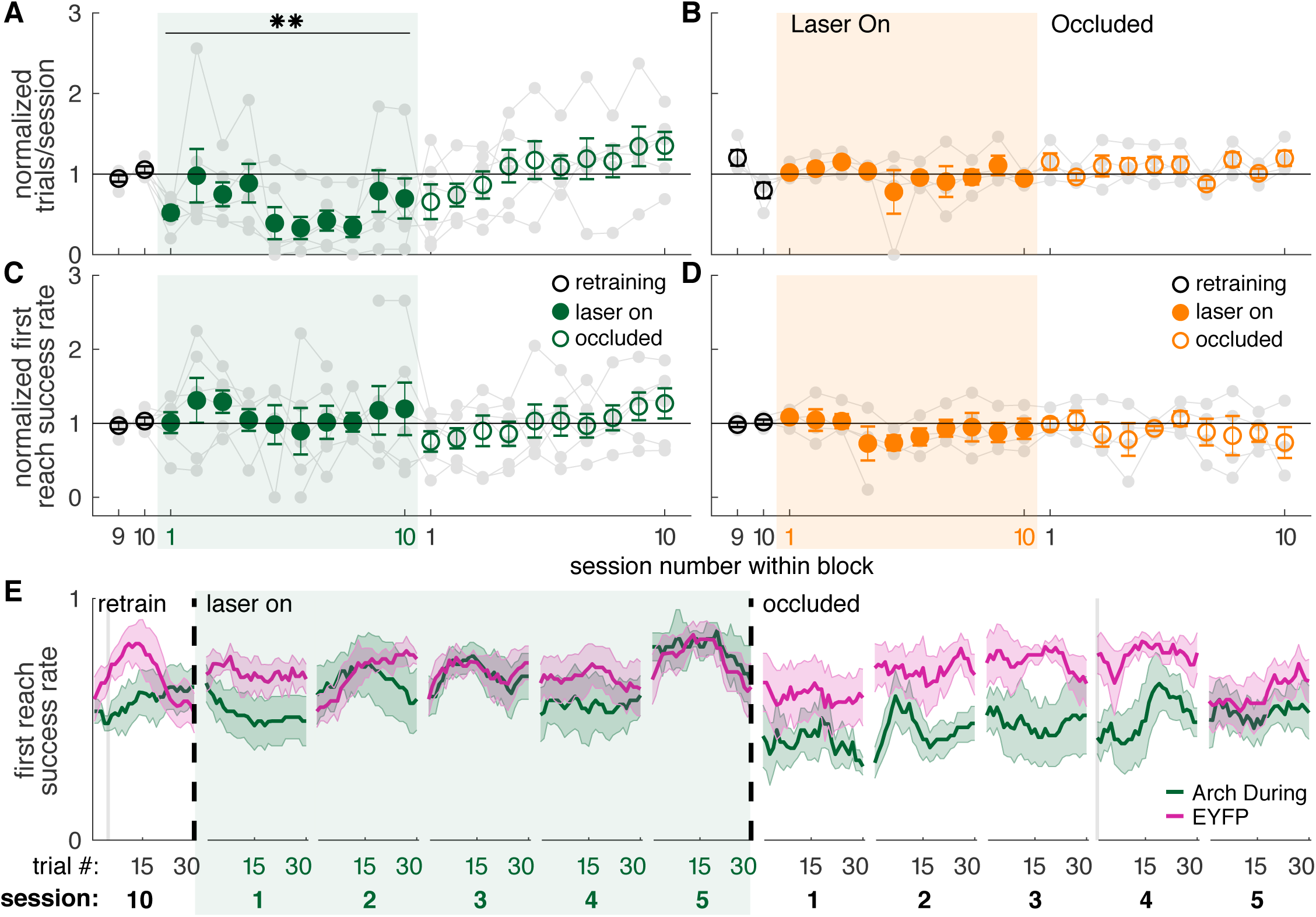
Dopamine neuron inhibition during reaches decreases the number of reaches performed but does not impair reach accuracy. **(A)** Average number of trials per session divided by the baseline number of trials for “during reach” inhibition. Grey lines represent individual rats. Linear mixed model: effect of laser: *t*(80) = −0.21, *P* = 0.84; interaction between laser and session: *t*(584) = −2.64, *P* = 8.47×10^-3^. **(B)** Same as (A) for “between reach” inhibition. Linear mixed model: effect of laser: *t*(80) = 0.93, *P* = 0.36; interaction between laser and session: *t*(584) = −1.52, *P* = 0.13. **(C)** Average first attempt success rate divided by baseline success rate for “during reach” inhibition. Linear mixed model: effect of laser: *t*(133) = 0.59, *P* = 0.56; interaction between laser and session: *t*(584) = −0.40, *P* = 0.69. **(D)** Same as (C) for “between reach” inhibition. Linear mixed model: effect of laser: *t*(133) = −0.64, *P* = 0.52; interaction between laser and session: *t*(584) = 0.10, *P* = 0.92. **(E)** Moving average of success rate across trials within individual sessions in the last retraining session, first 5 “laser on” sessions, and first 5 “occlusion” sessions. Shaded grey areas represent trials with a statistically significant difference between groups (Wilcoxon rank sum test, *P* < 0.01). Shaded colored areas in (E) and error bars in A-D represent s.e.m. Moving average of success rate within sessions for Arch Between rats is shown in Figure 4 – figure supplement 1. Data for individual Arch During and Arch Between rats are shown in Figure 4 – figure supplement 2. ** indicates p < 0.01 for the laser-session interaction term in panel A.

### Dopamine manipulations induce progressive changes in reach-to-grasp kinematics

The success rate analysis indicates that repeated dopaminergic stimulation progressively diminished reaching performance, but does not explain why performance worsened. To determine which aspects of reach kinematics were altered by dopaminergic manipulations, we used Deeplabcut to track individual digits, the paw, and the pellet (Figure 5; Bova et al., 2019, Mathis et al., 2018).

**Figure 5.**
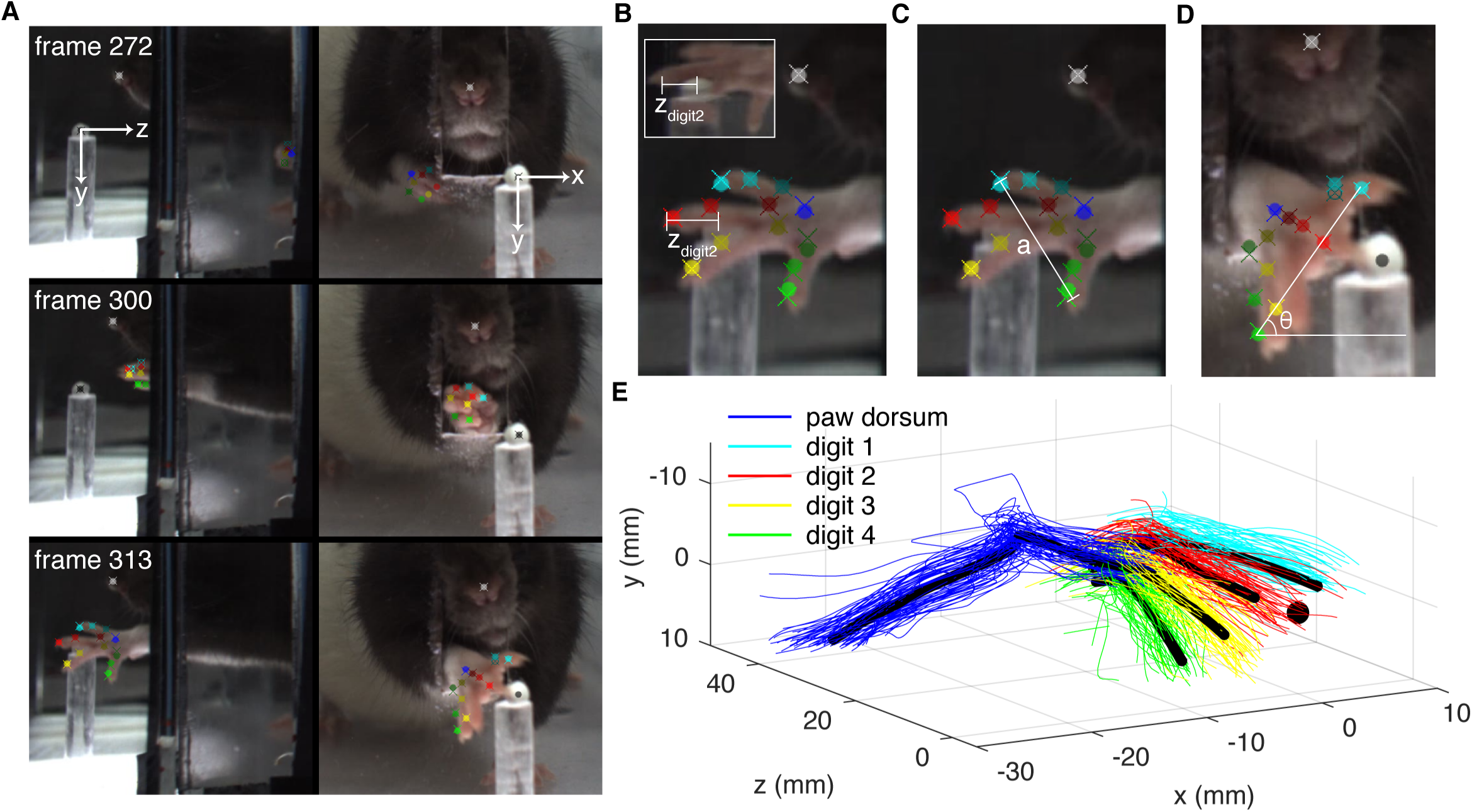
Paw and digit tracking with Deeplabcut. **(A)** Deeplabcut identification of digits, paw dorsum, nose, and pellet in individual video frames (side mirror and direct views). X, Y, and Z coordinates are in reference to the pellet. **(B)** Reach extent (z_digit2_) is the z-coordinate of the tip of the second digit. The end of a reach is defined as the moment z_digit2_ begins to increase (the digit tip moves back towards the box). Inset – mirror view of the palmar surface of the paw **(C)** Grasp aperture (a) is the Euclidian distance between the first and fourth digit tips. **(D)** Paw orientation is the angle (θ) between a line connecting the first and fourth digit tips and the floor. **(E)** Example 3-dimensional reconstruction of reaching trajectories from a single “retraining” session. Colored lines represent individual trials and black lines represent average trajectories of the paw dorsum and digit tips. Sugar pellet (black dot) is at (0,0,0).

Consistent with the success rate analysis, dopamine neuron stimulation during reaching caused progressive changes in reach-to-grasp kinematics. Reach extent (how far the paw extended in the direction of the pellet, z_digit2_) became progressively shorter with repeated stimulation during reaches (Figure 6A, Videos 1 and 2). This progressive change occurred both across and within sessions, and did not stabilize until the fifth session of dopamine neuron stimulation (Figure 6D and Figure 6 – figure supplement 2). Dopamine neuron stimulation during reaches also gradually narrowed grasp aperture at reach end (Figure 6E, F and Figure 6 – figure supplement 3), caused the paw to be more pronated at reach end (i.e., theta decreased) (Figure 6G, H and Figure 6 – figure supplement 4), and decreased the maximum reach velocity (Figure 6I, J and Figure 6 – figure supplement 5). Interestingly, kinematic measures continued to change even when success rate had plateaued (compare Figures 2E and 6). This is due to a “floor effect” for success rate – once the rat consistently misses the pellet, no further changes are detectable by this measure. When dopamine stimulation ceased (“occlusion” sessions), reach-to-grasp kinematics rapidly returned to baseline. As with success rate, there was individual variability in how quickly rats returned to pre-stimulation kinematics (Figures 6A, D-J and Figure 6 –figure supplements 2-5). All reach-to-grasp kinematics were unchanged in rats receiving dopamine neuron stimulation between reaches and EYFP control rats (Figures 6B-D, F, H, J and Figure 6 –figure supplements 1-5). In addition to histology, we verified opsin expression and fiber placement by performing “during reach” stimulation in rats previously stimulated between reaches. All rats showed kinematic changes with “during reach” stimulation not observed with “between reach” stimulation. This served as a positive control and reinforces the importance of the timing of dopamine neuron stimulation with respect to specific actions.

**Figure 6.**
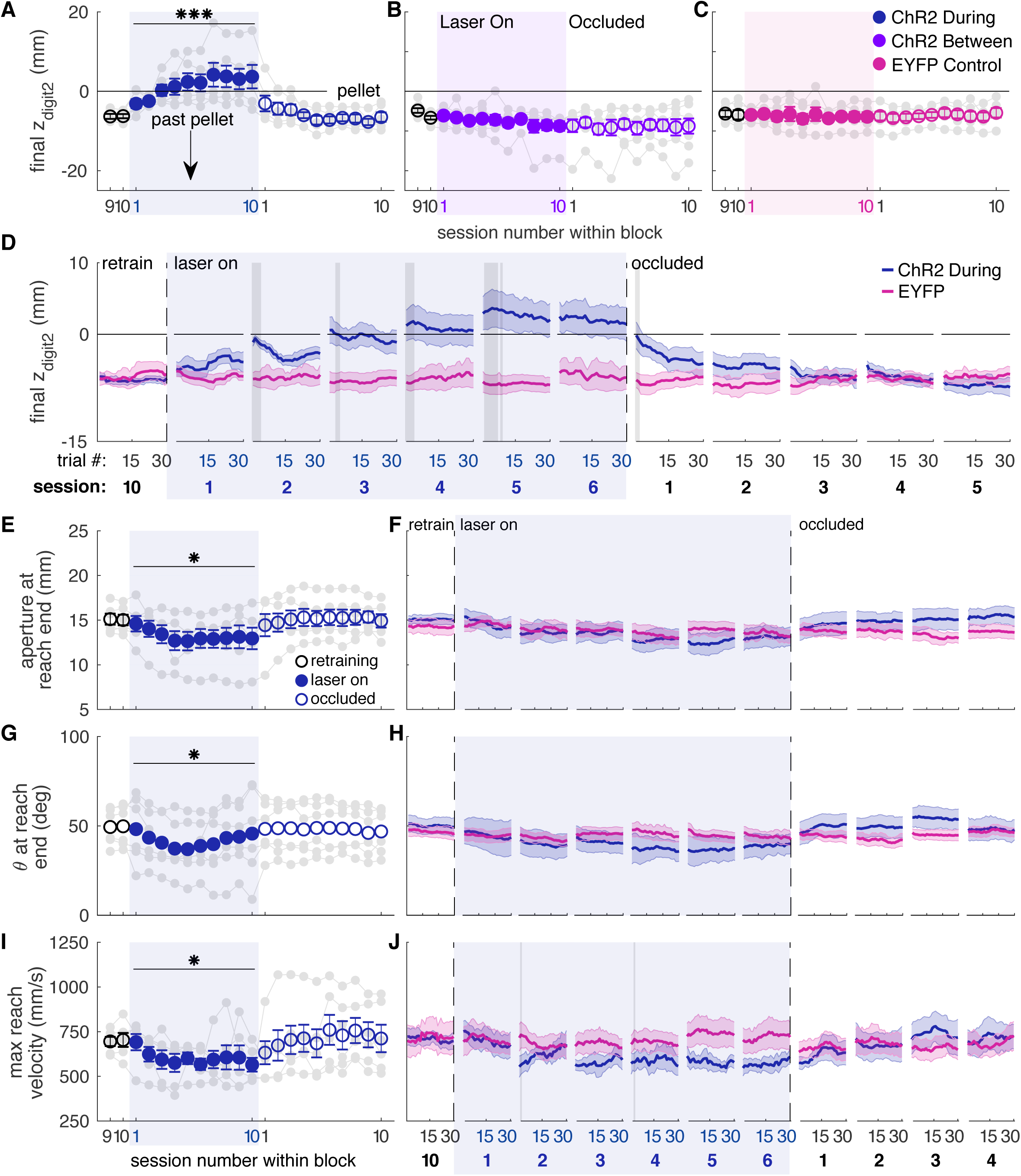
Dopamine neuron stimulation induces progressive changes in reach-to-grasp kinematics. **(A)** The average maximum reach extent progressively decreased across sessions with “during reach” stimulation. Linear mixed model: effect of laser: *t*(62) = 1.70, *P* = 0.09; interaction between laser and session: *t*(585) = 6.88, *P* = 1.59×10^-11^. Average maximum reach extent returned to baseline within the first “occlusion” session. Contrast testing (“retraining” session 10 vs. “occlusion” session 1): *t*(585) = 1.62, *P* = 0.11. **(B)** Same as (A) for “between reach” stimulation. Linear mixed model: effect of laser: *t*(62) = 0.02, *P* = 0.99; interaction between laser and session: *t*(585) = −0.43, *P* = 0.67. **(C)** Same as (A) and (B) for “during reach” illumination in control EYFP-injected rats. Linear mixed model: effect of laser: *t*(62) = 0.10, *P* = 0.92; interaction between laser and session: *t*(585) = −0.68, *P* = 0.50. **(D)** Moving average of maximum reach extent within the last “retraining” session, first 6 “laser on” sessions, and first 5 “occlusion” sessions. Grey shaded areas represent trials with a statistically significant difference between groups (Wilcoxon rank sum test, *P* < 0.01). **(E)** Average grasp aperture at reach end for “during reach” stimulation. Linear mixed model: effect of laser: *t*(48) = −1.34, *P* = 0.19; interaction between laser and session: *t*(585) = −2.19, *P* = 0.03. Average aperture returned to baseline within the first “occlusion” session. Contrast testing (“retraining” session 10 vs. “occlusion” session 1): *t*(585) = −0.87, *P* = 0.38. **(F)** Moving average of aperture at reach end within the last “retraining” session, first 6 “laser on” sessions, and first 4 “occlusion” sessions. **(G)** Same as (E) for paw orientation. Linear mixed model: effect of laser: *t*(74) = −2.52, *P* = 0.01; interaction between laser and session: *t*(585) = 0.19, *P* = 0.85. Average angle returned to baseline within the first “occlusion” session. Contrast testing (“retraining” session 10 vs. “occlusion” session 1): *t*(585) = 1.64, *P* = 0.10. **(H)** Moving average of paw angle at reach end across trials in the last (10th) “retraining” session, first 6 “laser on” sessions, and first 4 “occlusion” sessions. Grey shaded areas represent trials with a statistically significant difference between groups (Wilcoxon rank sum test, *P* < 0.01). **(I)** Same as (E) and (G) for maximum reach velocity. Linear mixed model: effect of laser: *t*(49) = −0.45, *P* = 0.65; interaction between laser and session: *t*(585) = −2.45, *P* = 0.01. Average velocity returned to baseline within the first “occlusion” session. Contrast testing (“retraining” session 10 vs. “occlusion” session 1): *t*(585) = −1.64, *P* = 0.10. **(J)** Moving average of maximum reach velocity within the last “retraining” session, first 6 “laser on” sessions, and first 4 “occlusion” sessions. Grey shaded areas represent trials with a statistically significant difference between groups (Wilcoxon rank sum test, *P* < 0.01). Shaded colored areas in D, F, H, J and error bars in A, B, C, E, G, I represent s.e.m. Similar data for ChR2 Between rats are shown in Figure 6 – figure supplement 1. Individual rat data are shown in Figure 6 – figure supplements 2-5. * indicates p < 0.05 for the laser or laser-session interaction terms in panels E, G, I. *** indicates p < 1.0 x 10^-10^ for the laser-session interaction term in panel A.

While dopamine neuron inhibition during reaching did not affect success rate (Figure 4C), it caused subtle changes in reach-to-grasp kinematics. Maximum reach extent lengthened slightly under dopamine neuron inhibition (that is, the paw extended further past the pellet, Figure 7A, C, Videos 3 and 4), in opposition to the effects of dopamine neuron stimulation. This effect almost reached significance in the linear mixed-effect model (p = 0.091, see Figure 7 caption), but a contrast test comparing laser day 10 to occlusion day 1 was significant (*t*(37) = −3.24, *P* = 0.003). Furthermore, reach extent consistently lengthened at the individual rat level (Figure 7A, gray markers) as well as across trials within sessions (Figure 7C, Figure 7 –figure supplement 2). Maximum reach velocity also decreased with dopamine inhibition (Figure 7H, I). This also was not quite significant in the linear mixed-effect model (p = 0.094, see Figure 7 caption), but there was a significant difference between laser day 10 and occlusion day 1 (contrast testing, *t*(33) = −2.49, *P* = 0.018). These data suggest that dopamine neuron stimulation and inhibition have roughly opposite effects on reach kinematics. Dopamine neuron inhibition did not significantly affect grasp aperture (Figure 7D, E) or paw orientation (Figure 7F, G), potentially due to ceiling effects. No kinematic changes were observed in rats that received dopamine neuron inhibition between reaches (Figure 7B and Figure 7 –figure supplements 1-5).

**Figure 7.**
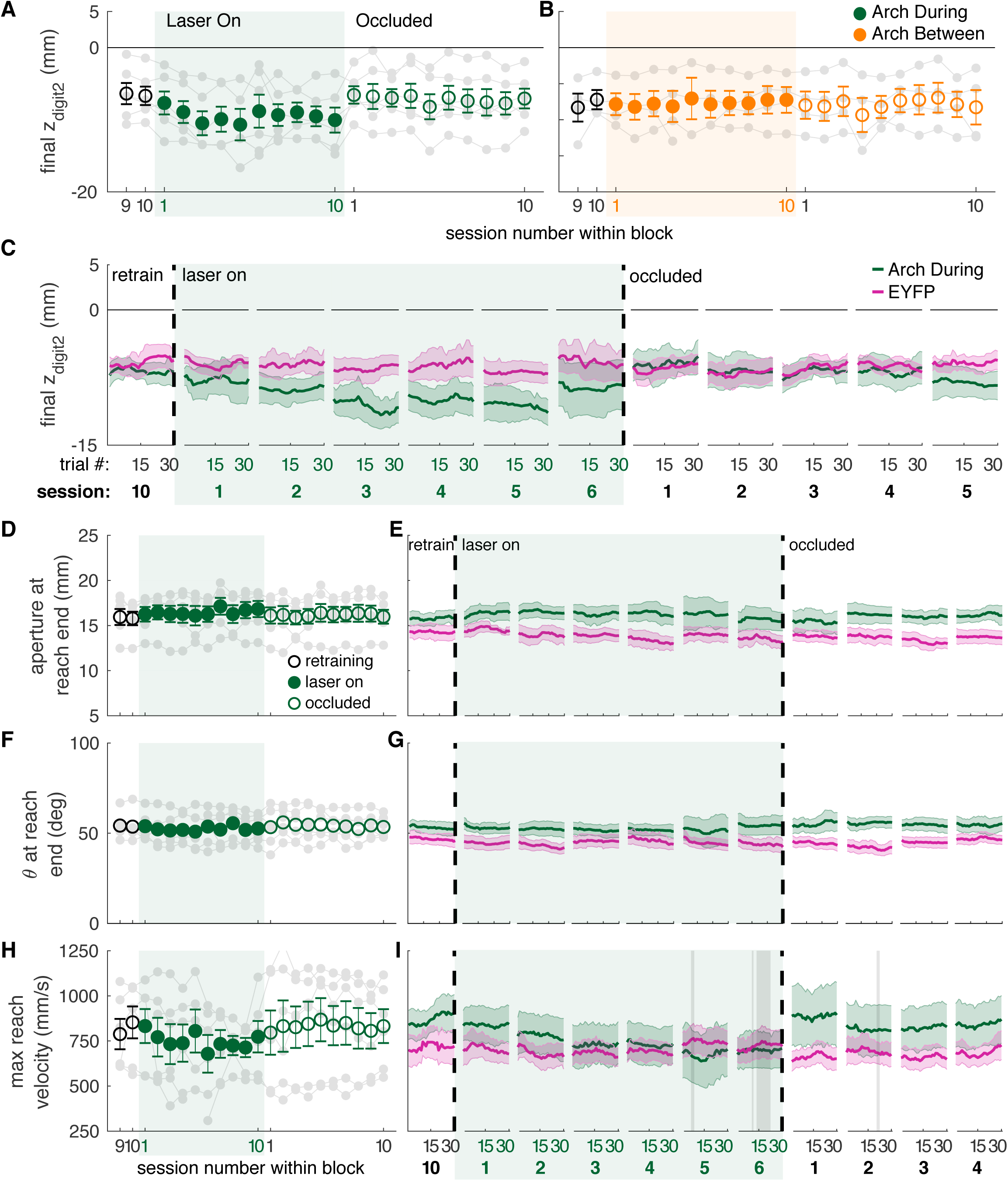
Dopamine neuron inhibition induces subtle changes in reach-to-grasp kinematics. **(A)** Average maximum reach extent across sessions for “during reach” inhibition. Linear mixed model: effect of laser: *t*(63) = −1.72, *P* = 0.09; interaction between laser and session: *t*(585) = 0.03, *P* = 0.98. **(B)** Same as (A) for “between reach” inhibition. Linear mixed model: effect of laser: *t*(63) = −0.23, *P* = 0.82; interaction between laser and session: *t*(585) = 0.99, *P* = 0.32. **(C)** Moving average of maximum reach extent within the last “retraining” sessions, first 6 “laser on” sessions, and first 5 “occlusion” sessions. **(D)** Same as (A) for aperture: effect of laser: *t*(48) = 0.53, *P* = 0.60; interaction between laser and session: *t*(585) = 1.76, *P* = 0.08. **(E)** Moving average of grasp aperture at reach end within the last “retraining” session, first 6 “laser on” sessions, and first 4 “occlusion” sessions. **(F)** Same as (A) and (D) for paw orientation: effect of laser: *t*(75) = −0.20, *P* = 0.84; interaction between laser and session: *t*(585) = −0.28, *P* = 0.78. **(G)** Moving average of paw angle at reach end within the last “retraining session, first 6 “laser on” sessions, and first 4 “occlusion” sessions. **(H)** Same as (A), (D), and (F) for maximum reach velocity: effect of laser: *t*(49) = −0.52, *P* = 0.60; interaction between laser and session: *t*(585) = −1.68, *P* = 0.09. **(I)** Moving average of maximum reach velocity within the last “retraining” session, first 6 “laser on” sessions, and first 4 “occlusion” sessions. Grey shaded areas represent trials with a statistically significant difference between groups (Wilcoxon rank sum test, *P* < 0.01). Shaded colored areas in C, E, G, I and error bars in A, B, D, F, H represent s.e.m. Similar data for Arch Between rats are shown in Figure 7 – figure supplement 1. Individual rat data are shown in Figure 7 – figure supplements 2-5.

### Dopamine manipulations disrupt reach-to-grasp coordination

Reach-to-grasp success requires precise coordination of a complex sequence of reach sub-movements. Reaches begin when the rat orients to the pellet with its nose, then lifts and aligns its paw at midline with the digits closed. As the forelimb advances towards the pellet, the digits extend and spread while the paw pronates. After the digits close to grasp the pellet, the forelimb and paw are raised and supinated to bring the pellet towards the mouth (Alaverdashvili & Whishaw, 2010, Whishaw et al., 2008, Whishaw & Pellis, 1990). Because fine motor coordination is impaired in patients with Parkinson Disease, including during reaching-to-grasp (Whishaw et al., 2002), we looked to see if the coordination of reach sub-movements was affected by dopamine neuron stimulation or inhibition.

Dopamine neuron stimulation during reaching altered the coordination of digit spread (aperture) and paw pronation (orientation) with respect to paw advancement (Figures 8 and 9). Aperture increased earlier (when the paw was further from the pellet) in “during reach” stimulation sessions compared to “retraining” or “occlusion” sessions (Figure 9A, B). Thus, during dopamine stimulation, aperture was smaller at reach end but larger (on average) at matched distances from the pellet (Figures 6E, 8B, 9C). “During reach” dopamine neuron inhibition had the opposite effect – paw aperture began to increase when the paw was closer to the pellet compared to “retraining” or “occlusion” sessions (Figures 9A, B). Similar changes occurred with paw orientation: during sessions with dopamine neuron stimulation, paw pronation began further from the pellet (Figures 9D, E, F). Dopamine neuron inhibition, however, did not affect the relationship between paw orientation and paw advancement. As for other kinematic changes, the changes in coordination progressed across sessions (most evident in Figures 9C, F). No changes were observed in rats that received dopamine stimulation or inhibition between reaches or in EYFP control rats (Figures 9B, C, E, F and Figure 9 – figure supplements 1-4). Together, these results suggest that dopamine neuron stimulation accelerates transitions between reach sub-movements, while dopamine neuron inhibition has the opposite effect.

**Figure 8.**
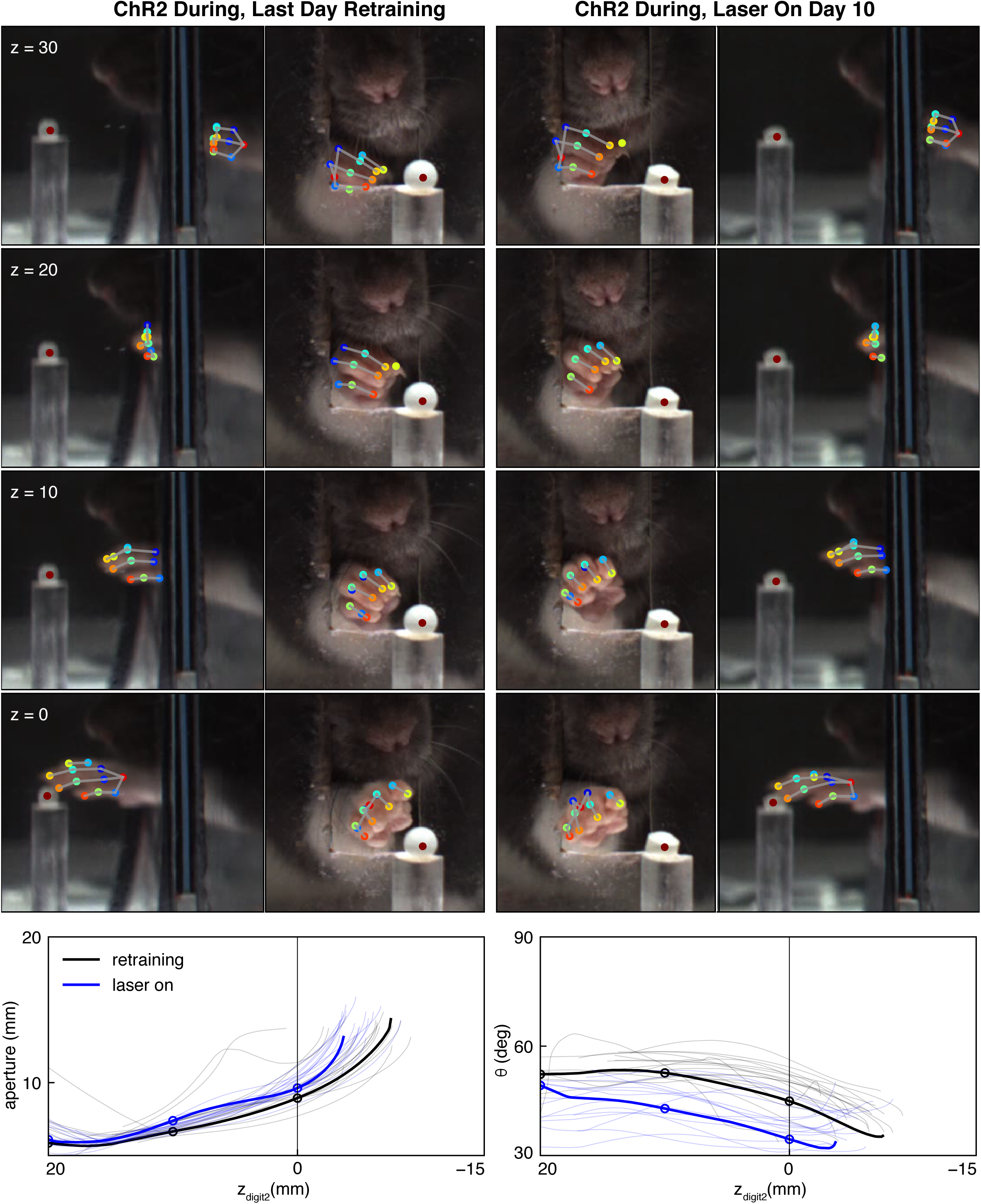
Dopamine neuron stimulation alters the coordination between digit movements and paw advancement. **(A)** Sample frames from single reaches at the end of “retraining” and “laser on” sessions from the same rat. Outer columns show the mirror views corresponding to the direct camera views in the inside columns. After 10 days of “during reach” stimulation, the rat pronates its paw and spreads its digits further from the pellet as the paw advances. **(B)** Aperture as a function of the z-coordinate of the second digit tip. Solid black and blue lines correspond to the reaches shown in (A). Thin black and blue lines are the traces for other reaches in the same sessions. Circles indicate apertures at the corresponding z_digit2_ values in A. **(C)** Same as (B) but for paw orientation.

**Figure 9.**
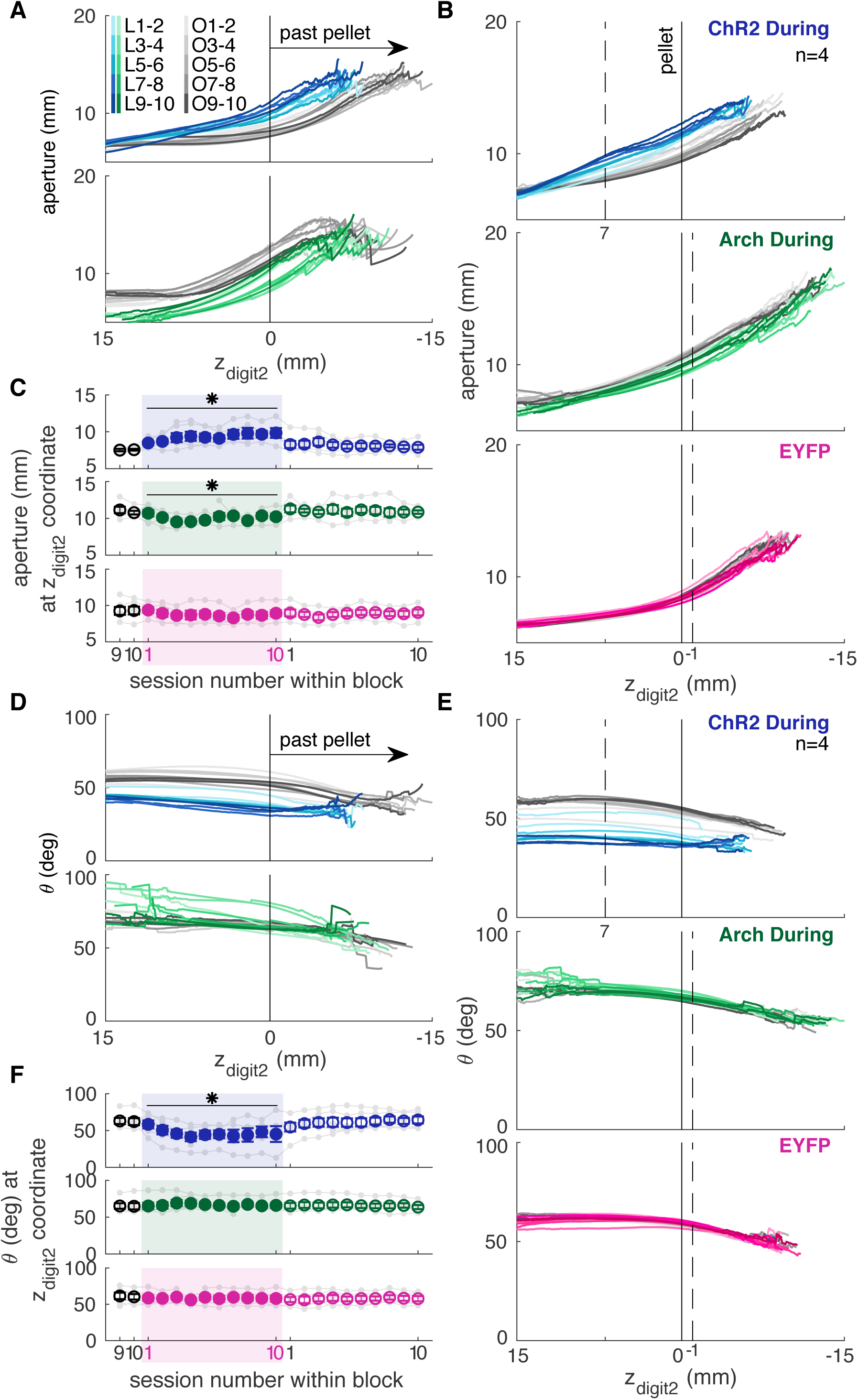
Dopamine neuron manipulations disrupt coordination of reach-to-grasp movements. **(A)** Mean aperture as a function of paw advancement (z_digit2_, pellet at z_digit2_=0) across “laser on” and “occlusion” sessions for exemplar rats. All rats are shown in Figure 9 – figure supplement 3. L1-2, O1-2, … indicate laser on sessions 1-2, occlusion sessions 1-2, etc. **(B)** Mean aperture as a function of paw advancement across “laser on” and “occlusion” sessions averaged across all rats. 4 of 6 “ChR2 During” rats are included because 2 rats’ reaches were too short in several sessions to produce a meaningful average (the average for all 6 ChR2 During rats, ChR2 Between rats, and Arch Between rats are shown in Figure 9 – figure supplement 1). All rats were included for other groups. Dashed lines indicate the z_digit2_ coordinate where data are sampled in (C) for each group. A more proximal z_digit2_ was chosen for “ChR2 During” because the majority of “laser on” reaches for this group did not extend past z_digit2_ = −1 mm. **(C)** Average grasp aperture at the z_digit2_ coordinates indicated by the dashed lines in (B) across sessions. “During reach” stimulation gradually increased aperture at 7 mm from the pellet (linear mixed model including all 6 “during reach” rats: effect of laser: *t*(607) = 2.39, *P* = 0.02; interaction between laser and session: *t*(607) = 2.40, *P* = 0.02). “During reach” inhibition decreased aperture at 1 mm past the pellet (linear mixed model: effect of laser: *t*(607) = −2.04, *P* = 0.04; interaction between laser and session: *t*(607) = 0.67, *P* = 0.51). SNc illumination in EYFP-injected rats had no effect on aperture at 1 mm past the pellet (linear mixed model: effect of laser: *t*(607) = −0.57, *P* = 0.57; interaction between laser and session: *t*(607) = −0.61, *P* = 0.54). Grey points indicate data from individual rats. **(D)** Mean paw orientation as a function of paw advancement towards the pellet across “laser on” and “occlusion” sessions for exemplar rats. All rats are shown in Figure 9 – figure supplement 4. **(E)** Mean paw orientation as a function of paw advancement across “laser on” and “occlusion” sessions averaged across rats. Dashed lines indicate z_digit2_ coordinates where data are sampled in (F) for each group. 4 of 6 “ChR2 During” rats are included because 2 rats’ reaches were too short in several sessions to produce a meaningful average (the average for all 6 ChR2 During rats, ChR2 Between rats, and Arch Between rats are shown in Figure 9 – figure supplement 2). **(F)** Average paw orientation at z_digit2_ coordinates indicated by dashed lines in (E) across all sessions. “During reach” stimulation caused a gradual increase in pronation (i.e., a smaller angle) at 7 mm from the pellet (linear mixed model including all 6 “during reach” rats: effect of laser: *t*(607) = −2.34, *P* = 0.02; interaction between laser and session: *t*(607) = −2.33, *P* = 0.02). “During reach” inhibition had no effect on paw orientation at 1 mm past the pellet (linear mixed model: effect of laser: *t*(607) = 0.88, *P* = 0.38; interaction between laser and session: *t*(607) = −0.55, *P* = 0.58). SNc illumination in EYFP-injected rats had no effect on paw orientation at 1 mm past the pellet (linear mixed model: effect of laser: *t*(607) = −0.51, *P* = 0.61; interaction between laser and session: *t*(607) = 0.31, *P* = 0.76). Grey points indicate data from individual rats. * indicates p < 0.05 for either the laser or laser-session interaction terms in panels C and F.

### Dopamine neuron stimulation establishes distinct reach-to-grasp representations

Dopamine neuron stimulation gradually induced changes in reach-to-grasp kinematics, but kinematics rapidly recovered to baseline when the laser was occluded. We next asked if reinstating dopamine neuron stimulation would again gradually alter reach kinematics. Following testing with the laser occluded, six ChR2-injected rats that had received “between reach” stimulation performed additional “during reach” stimulation sessions. These continued until reach-to-grasp kinematics were impaired (average: 3.17 ± 0.98 sessions). Once kinematics were impaired, rats performed an additional one or two 30-minute sessions during which the laser alternated every 5 trials between being off and on during reaches (Figure 10A).

**Figure 10.**
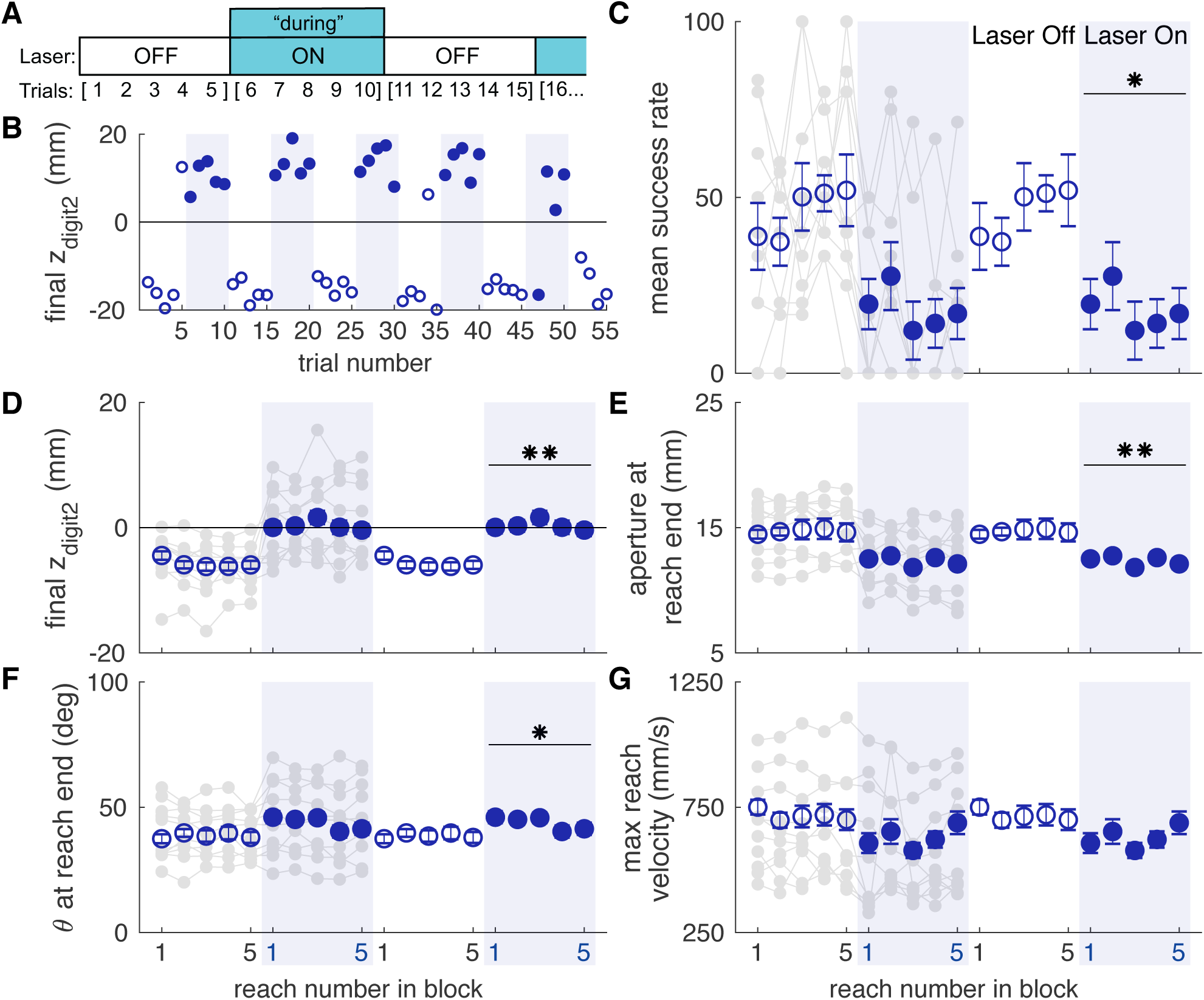
Dopamine neuron stimulation induces distinct reach-to-grasp kinematics that depend on current dopamine stimulation. **(A)** Schematic of alternating stimulation test sessions. **(B)** Example session from one rat with maximum reach extent plotted for every trial. Some blocks have fewer than 5 trials if the rat did not reach for the pellet after breaking the IR beam. **(C)** Average first attempt success rate during “laser off” and “laser on” blocks. Data are repeated to show “off to on” and “on to off” transitions. Grey lines show individual rat data. Linear mixed model: effect of laser: *t*(78) = −0.50, *P* = 0.62; interaction between laser and trial within block: *t*(78) = −2.35, *P* = 0.02. **(D)** Average maximum reach extent during “laser off” and “laser on” blocks. Linear mixed model: effect of laser: *t*(78) = 2.70, *P* = 8.47×10^-3^; interaction between laser and trial within block: *t*(78) = 1.32, *P* = 0.19. **(E)** Average aperture at reach end across “laser off” and “laser on” blocks. Linear mixed model: effect of laser: *t*(78) = −2.83, *P* = 5.92×10^-3^; interaction between laser and trial within block: *t*(78) = −0.79, *P* = 0.43. **(F)** Average paw orientation at reach end across “laser off” and “laser on” blocks. Linear mixed model: effect of laser: *t*(78) = 2.57, *P* = 0.01; interaction between laser and trial within block: *t*(78) = −0.34, *P* = 0.73. **(G)** Average maximum reach velocity across “laser off” and “laser on” blocks. Linear mixed model: effect of laser: *t*(78) = −1.24, *P* = 0.22; interaction between laser and trial within block: *t*(78) = 0.01, *P* = 0.99. * indicates p < 0.05 for effect of laser in panel F. ** indicates p < 0.01 for effect of laser in panel D.

Rats transitioned rapidly between “normal” and “impaired” reach-to-grasp kinematics with the laser off and on, respectively. The mean success rate dropped within a single trial of laser stimulation and improved within one trial when laser stimulation was removed (Figure 10C, Video 5). Similarly, reach kinematics required only one trial to switch between normal and aberrant reaching patterns. Laser stimulation at the beginning of an “on” block (trial 1) caused immediate decreases in maximum reach extent and digit aperture, which remained steady for the remaining “Laser On” trials. Similarly, maximum reach extent and digit aperture immediately increased upon cessation of dopamine neuron stimulation (trial 1, “Laser Off”) and remained steady throughout the “Laser Off” block (Figure 10D, E). There was also a significant change in paw orientation at reach end with dopamine neuron stimulation (Figure 10F). However, pronation decreased in these rats unlike in the “ChR2 During” group (Figure 6G). There was no significant difference in maximum reach velocity between “Laser On” and “Laser Off” blocks (Figure 10G). These data indicate that once distinct reaching kinematics have been established by repeated dopaminergic manipulations, current reach kinematics are determined by the activity of nigral dopamine neurons on that trial.

### Dopamine neuron stimulation induces context- and history-dependent abnormal involuntary movements

To verify fiber placement and opsin expression prior to reaching experiments, we placed rats in a clear cylinder and illuminated SNc with blue light of varying intensity (Figure 11A). We predicted that rats with well-placed fibers expressing high levels of ChR2 would develop increasingly worse abnormal involuntary movements (AIMs) as laser intensity increased. To our surprise, rats that subsequently developed markedly abnormal reach kinematics during the skilled reaching task appeared unaffected by dopamine neuron stimulation in the cylinder (AIMs Test 1, Figure 11B-F). In “post-reaching” cylinder sessions (AIMS Test 2, Figure 11B), however, dopamine neuron stimulation elicited markedly abnormal movements (Figure 11C-E, Figure 11, Video 6). Furthermore, while AIMs were obvious in the context of the cylinder, the same (or higher) stimulation intensities delivered while rats were reaching failed to elicit abnormal movements (other than altered reach kinematics). Thus, the expression of dopamine-dependent AIMs depends not only on current levels of dopamine neuron activation, but the history of prior activation and the current behavioral context.

**Figure 11.**
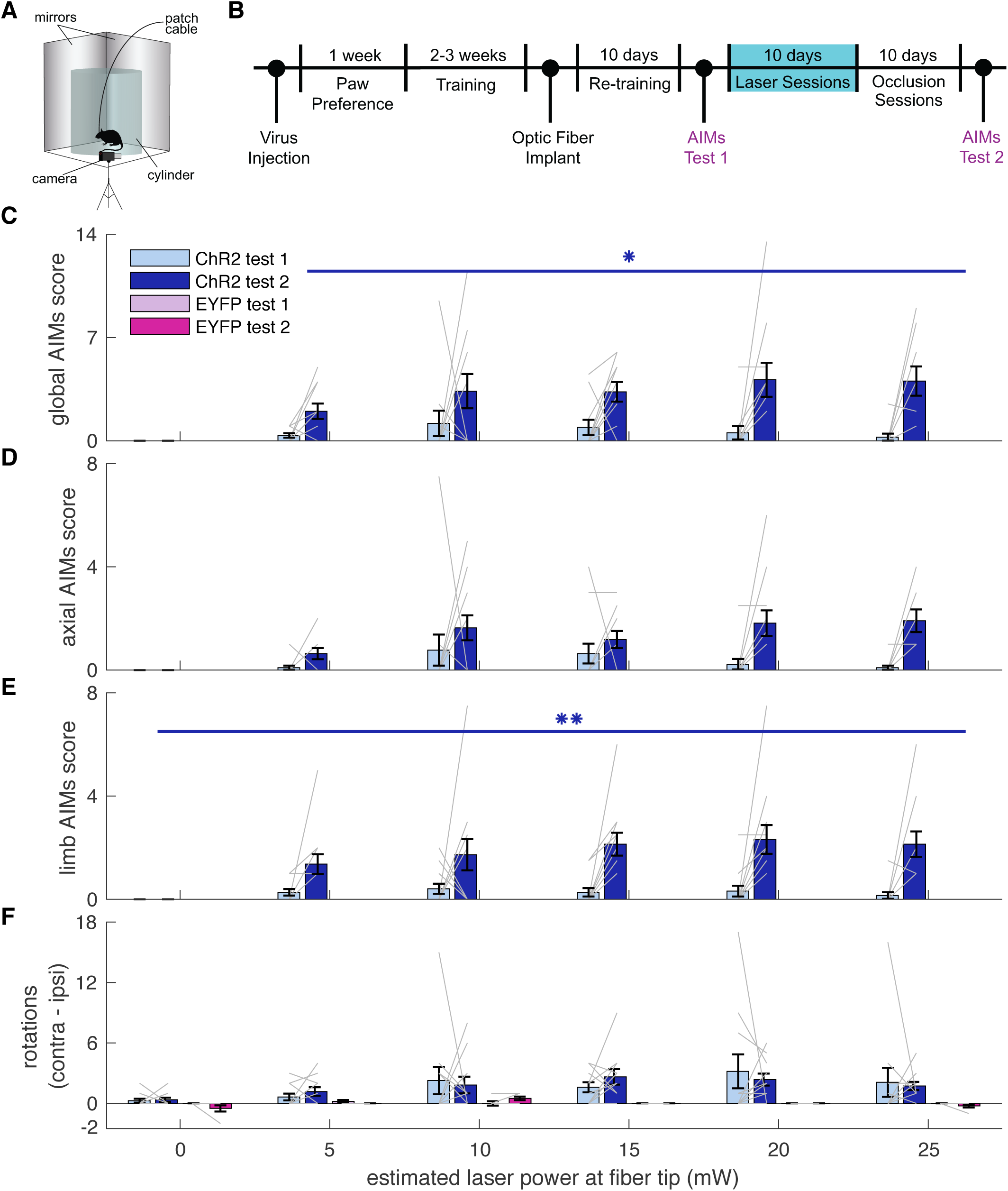
Dopamine neuron stimulation induces context- and history-dependent abnormal involuntary movements. **(A)** Experimental set-up for AIMs test. **(B)** Timeline of experiment. The first AIMs test took place one day before skilled reaching testing; the second took place one day after the last “occlusion” session. Repeated from Figure 1A for convenience. **(C)** Average global AIMs scores vs. estimated power at the fiber tip. Global AIMs increased with increasing laser power and from test day 1 to 2 in ChR2-injected rats (linear mixed model: interaction between test number and laser power: *t*(164) = 2.57, *P* = 0.01). EYFP-injected rats did not display AIMs (linear mixed model: interaction between test number and laser power: *t*(164) = 0.00, *P* = 1.00). Gray lines represent data from individual rats. Error bars represent s.e.m. across animals. **(D)** Average axial AIMs scores. ChR2: linear mixed model: interaction between test number and laser power: *t*(165) = 1.91, *P* = 0.06. EYFP: *t*(165) = 0.00, *P* = 1.00. **(E)** Average limb AIMs scores. A linear mixed-effects model found a significant interaction between test number and laser power in ChR2-injected rats: *t*(164) = 2.81, *P* = 5.51×10^-3^. EYFP-injected rats did not display limb AIMs: *t*(164) = 0.00, *P* = 1.00. **(F)** Difference between average number of contralateral and ipsilateral (relative to hemisphere implanted with optical fiber) rotations. A positive score indicates a bias towards contralateral spins and a negative score indicates a bias towards ipsilateral spins. ChR2-injected rats did not increase the number of contralateral spins between test 1 and test 2, nor did laser power affect rotational behavior. Linear mixed model: interaction between test number and laser power: *t*(164) = −0.39, *P* = 0.69. EYFP-injected rats did not show a bias in either direction with laser stimulation: *t*(164) = 0.10, *P* = 0.92. (* P < 0.05, ** P < 0.01 for ChR2-injected rats).

## Discussion

Our goal was to determine how midbrain dopamine neuron manipulations affect dexterous skill. Our results revealed a role for dopamine in motor learning, as repeated dopamine manipulations induced gradual changes in reach-to-grasp kinematics. These manipulations not only affected gross performance measures (e.g., velocity and amplitude) but also disrupted coordinated execution of reach sub-movements. Once dopamine stimulation-induced changes were established, reach-to-grasp kinematics depended strongly on the dopamine status of the current trial. Furthermore, these effects were temporally specific – only manipulations during reaches influenced forelimb kinematics. Finally, the effect of dopamine on motor control is context-dependent, as the same dopamine stimulation in different situations induced distinct behavioral responses.

The history-dependent effects of dopamine on skilled reaching are superficially consistent with reinforcement learning models (Schultz, 2019). While most evidence for dopamine signals encoding RPEs comes from paradigms in which animals choose between discrete actions (e.g., press a right or left lever), recent studies suggest that dopamine encodes RPE-like “performance prediction errors” for complex behaviors with greater degrees of freedom (Beeler et al., 2010, Gadagkar et al., 2016). It is plausible that dopamine neuron excitation/inhibition creates an artificially reinforcing/discouraging signal that influences subsequent reaches. Within this framework, dopamine neuron stimulation (or inhibition), regardless of reach outcome, should gradually alter reach kinematics. Furthermore, since there are many possible failure mechanisms, the changes in kinematics should be unpredictable. However, the effects of dopamine manipulations on reach kinematics were consistent, with dopamine neuron stimulation and inhibition inducing essentially opposite changes. This suggests that kinematic changes do not result purely from performance prediction error signals, but that dopamine intrinsically biases movement kinematics in a consistent direction.

Motion tracking data provide insight into the nature of this intrinsic dopamine bias. A common interpretation of dopamine’s role in movement is that it regulates “vigor,” which has been defined as the speed, frequency, and amplitude of movements (Dudman & Krakauer, 2016). Our dopamine manipulations influenced “vigor” in unexpected ways: dopamine neuron stimulation decreased, and inhibition increased, movement amplitude (reach extent). Furthermore, both stimulation and inhibition decreased movement speed. These effects apparently contradict previous work directly correlating dopaminergic tone with movement velocity and/or amplitude (Carr & White, 1987, Leventhal et al., 2014, Panigrahi et al., 2015).

This discrepancy may be due to different demands on the motor system. “Vigor” assays generally demand movement along one dimension. For example, mice manipulating a joystick (Panigrahi et al., 2015), or humans moving a manipulandum to a target (Baraduc et al., 2013, Mazzoni et al., 2007) make forelimb/arm movements across large joints more or less along a single vector. In such tasks, dopamine-depleted subjects consistently make hypometric, bradykinetic movements. In contrast, skilled reaching comprises a sequence of precisely-coordinated submovements (Klein & Dunnett, 2012). Stimulation caused paw pronation and digit spread to occur earlier along the reach trajectory (farther from the pellet), while inhibition delayed these submovements with respect to paw extension. This is consistent with evidence that dopamine regulates initiation of/transitions between movements (da Silva et al., 2018). That is, dopamine may increase the probability of initiating the next submovement in the skilled reaching sequence. This may be interpreted in a “vigor” framework as striatal dopamine invigorating the next submovement at the expense of the current one. In a complex, multi-component movement like skilled reaching, this would cause premature transitions and compress the overall reach-to-grasp sequence.

While kinematic changes developed gradually across sessions, once established they depended on the dopamine status of the current trial (Figure 10). This suggests that dopamine stimulation instantiated distinct representations of movement kinematics, which were selected for execution by current dopamine neuron activity. These representations could be stored in any motor-related brain region, or as an emergent property of larger motor circuits. However, the fact that we stimulated preferentially over SNc suggests that they are stored in striatum. Consistent with this idea, recent work identified subpopulations of direct pathway medium spiny neurons associated with dyskinesias after levodopa treatment in dopamine-depleted mice (Ryan et al., 2018). Furthermore, specific activation of these direct pathway MSNs induced dyskinesias (Girasole et al., 2018). These results suggest that subpopulations of striatal output neurons encode specific movement kinematics that are sensitive to striatal dopamine levels.

The abrupt transitions between aberrant and baseline reach kinematics are reminiscent of “on/off” motor fluctuations observed in people with PD. With disease progression and prolonged treatment, patients often display sudden transitions between severe bradykinesia, good motor control, and levodopa-induced dyskinesias (Chou et al., 2018). Because disease duration, degree of dopamine loss, and magnitude of treatment-related dopamine fluctuations are correlated (Abercrombie et al., 1990, de la Fuente-Fernández et al., 2004), the root cause of motor fluctuations in PD patients is difficult to identify. Our results indicate that large, temporally specific dopamine fluctuations are sufficient to cause dramatic dopamine-dependent changes in movement kinematics, even in otherwise healthy subjects. This suggests that large swings in striatal dopamine are sufficient to generate motor fluctuations, independent of the degree of dopamine denervation.

The motor effects of dopamine neuron stimulation also depended on behavioral context. Dopamine neuron stimulation had almost no effect on stimulation-naïve rats in a clear cylinder. Rats engaged in skilled reaching during dopamine neuron stimulation continued to engage in the task, with few abnormal involuntary movements during reaching. However, the same stimulation parameters delivered to previously-stimulated rats in clear cylinders induced markedly abnormal limb and body movements (Figure 11 and Video 6). Broadly, this is consistent with the idea that dopamine regulates the “vigor” of movements selected based on the current behavioral context (Yttri & Dudman, 2016). That is, in the reaching chamber, rats approach the reaching slot to perform a (dopamine-modified) reach because that is the appropriate action in that context. Conversely, with no specific goal-directed actions suggested by the cylinder context, dopamine equally invigorates many potential movements. This leads to seemingly random abnormal involuntary movements (Bastide et al., 2015). Interestingly, the severity of experimental levodopa-induced dyskinesias depends on behavioral context (Lane et al., 2011). Finally, this context dependence of dopaminergic effects on motor control has parallels in clinical phenomenology: people with PD often can perform goal-directed movements despite the presence of significant levodopa-induced dyskinesias.

There are several limitations of this study. First, we did not record from dopamine neurons or measure dopamine release during optogenetic manipulations. It is therefore not clear how striatal dopamine levels were altered relative to normal reach-related dopamine dynamics, or if repeated stimulation changed spontaneous or optically evoked dopamine release (Saunders et al., 2018). Given the relatively high optical stimulation power (20 mW at the fiber tip) and frequency (20 Hz) used, we suspect that we induced supraphysiologic dopamine release (Patriarchi et al., 2018). Nonetheless, supra/infraphysiologic manipulations (e.g., lesion studies) can provide important insights into normal function. Furthermore, supraphysiologic dopamine fluctuations are relevant to pathologic states like PD, in which striatal dopamine can transition over minutes to hours between very low and high levels (Abercrombie et al., 1990, de la Fuente-Fernández et al., 2004). Second, we stimulated over SNc. It is therefore unclear how ventral tegmental area (VTA) stimulation would influence skilled reaching, and whether stimulating specific nigral projection fields (e.g., striatal subregions or motor cortex, Guo et al., 2015, Hosp et al., 2011, Zhang et al., 2017) would differentially affect reach kinematics. Finally, while we found that dopamine neuron manipulations during, but not between, reaches affected reach kinematics, the timing of when dopamine manipulations exert their effects could be parsed more precisely. Our “during reach” timing covered approach to the pellet, the reach itself, and immediately after the grasp during pellet consumption. Activation of different terminal fields at different times with respect to behavior may have dissociable effects on task performance.

In summary, temporally specific dopamine signals cause gradual changes in dexterous skill performance separable from pure “vigor” effects. These changes are durable, and expressed in a dopamine-dependent manner on a reach-by-reach basis. This phenomenon has clinical analogy with rapid motor fluctuations in PD patients. It may, therefore, serve as a useful paradigm in which to study the underlying neurobiology of motor fluctuations in PD, as well as address fundamental questions regarding how dopamine and basal ganglia circuits regulate skilled movements.

## Materials and methods

### Rats

All animal procedures were approved by the University of Michigan Institutional Animal Care & Use Committee. Numbers of rats included in each experimental group and analysis are indicated in figure legends and the main text. Male (n = 23) and female (n = 15) tyrosine hydroxylase-Cre^+^ (TH-Cre^+^) rats were housed in groups of 2-3 on a reverse light/dark cycle prior to optical fiber implantation. Following surgery, rats were housed individually to protect the implant. All testing was carried out during the dark phase. Food restriction was imposed on all animals during the training and testing periods for no more than 6 days in a row such that rats’ weights were kept ∼85-90% of their free-feeding weight. Water was available *ad libitum* in their home cages. Eight rats were excluded from the analysis due to either poor opsin expression or misplaced optical fibers (number of rats excluded: Group 1: n = 1; Group 2: n = 3; Group 3: n = 3; Group 4: n = 0; Group 5: n = 1). Judgment on whether to include subjects was made by investigators blinded to experimental groups and outcomes.

### Stereotaxic surgeries

Before pre-training for skilled reaching, rats were anesthetized with isoflourane (5% induction and 2-3% maintenance) and bilaterally injected in the SNc (M-L ± 1.8 mm; A-P −5.2 mm, –6.2 mm; D-V –7.0 mm, −8.0 mm) with AAV-EF1*α*-DIO-hChR2(H134R)-EFYP, AAV-EF1*α*-DIO-eArch3.0-EYFP, or AAV-EF1*α*-DIO-EYFP (UNC vector core). 1 μl of virus (titer: 3.4-4.2×10^12^ vg/ml) was injected per site (4 ul total per hemisphere) at a rate of 0.1 μl/min. After reaching stable performance on the skilled reaching task, optical fibers (multimode 200 μm core, 0.39 NA, Thor Labs FT200EMT) embedded in stainless steel ferrules (2.5 mm outer diameter, 230 μm bore size, Thor Labs #SF230-10) were implanted above SNc contralateral to the rat’s preferred reaching paw (M-L ± 2.4 mm, A-P −5.3 mm, D-V −7.0 mm). Optical fibers were calibrated before implantation to determine optical power at the fiber tip as a function of laser output power, which was continuously monitored during experiments by “picking off” 10% of the laser output with a beamsplitter. Rats recovered for at least 7 days after surgical procedures before beginning behavioral training or testing.

### Skilled reaching

#### Automated reaching system

Training and testing were carried out in custom-built skilled reaching chambers housed within soundproof, ventilated cabinets (Figure 1D, Bova et al., 2019, Ellens et al., 2016). Infrared sensors (HoneyWell, Morriston, NJ) were aligned so that the beam was directed through the back of the chamber. A reaching slot (1.1 x 7 cm) was cut into the front panel of the chamber 3.5 cm from the floor. One mirror was placed on either side of the front reaching chamber and angled to allow side views of the paw during reaches. A linear actuator with three position digital control (Creative Werks Inc., Des Moines, IA) and connected to an acrylic pellet delivery rod was mounted in a custom frame below the support box. The pellet delivery rod extended through a funnel mounted to the top of the frame. Before each session, the actuator was positioned so that the delivery rod was aligned with the right or left edge of the slot according to each rat’s paw preference 15 mm from the front of the reaching slot.

Videos were recorded at 300 frames-per-second and 2400 x 1024 pixels by a high-definition color digital camera (acA2000-340kc, Basler, Ahrensburg, Germany) mounted in front of the reaching slot. A camera-link field programmable gate array (FPGA) frame-grabber card (PCIe 1473R, National Instruments, Austin, TX) acquired the images, and an FPGA data acquisition (DAQ) task control card (NI PCIe 7841R) provided an interface with the behavior chamber and optogenetic system. The real-time FPGA card detected pixel intensity changes within a “region of interest” in front of the reaching slot visible in the side mirror views (Figure 1D), allowing videos of the reaching event (“video trigger”) to be captured. 300 frames pre-trigger and 1000 frames post-trigger were saved. A second camcorder was placed above the reaching chamber to record the entire session at 60 frames-per-second (HC-V110, Panasonic).

#### Trial performance

Custom LabVIEW software controls the experiment (Bova et al., 2019, Ellens et al., 2016). Each training session begins with the pellet delivery rod in the “ready” position - halfway between the bottom of the reaching chamber and the reaching slot. When the rat breaks the IR beam at the back of the chamber, the pellet delivery rod rises to the bottom of the reaching slot. When the reaching paw passes the front plane of the chamber into the “region of interest” and surpasses the minimum threshold of pixel intensity, video acquisition is triggered, time-stamped, and labeled with the trial number. Two seconds after the video is triggered, the pellet delivery arm lowers into the pellet funnel to pick up a new pellet and then resets to the “ready” position, allowing the rat to initiate a new trial.

#### Pre-training

“Pre-training” consists of familiarizing the rats with the reaching chamber, evaluating them for paw-preference, training them to reach for the linear actuator, and training them to request a pellet by moving to the back of the chamber. A week before pre-training, rats were placed on food restriction and introduced to the sucrose reward pellets in their home cages. On day 1 of pre-training, piles of five pellets each were placed in the front and rear of the skilled reaching chamber to encourage exploration of the entire chamber. Once rats ate these pellets, they were evaluated for paw-preference.

Rats were allowed to eat 3 pellets (held in forceps through the reaching slot) with their tongues. The experimenter then began to pull the pellet away from the rat so that it could not be obtained by licking. Therefore, the rat was forced to reach with its paw to retrieve the pellet. Paw preference was assigned to the paw used for the majority of the first eleven reaches. Once paw preference was determined, animals were trained to reach for the pellet delivery rod. As the rat reached, the experimenter pulled the forceps back so that the rat’s paw would extend to a pellet on the delivery rod. Once rats reached for the delivery rod 10 times without being baited by the experimenter, they began training to request pellets.

Rats began training in the center of the chamber with the pellet delivery rod set to the “ready” position. The experimenter placed a pellet in the rear of the chamber to bait the rat to break the rear IR beam, causing the delivery rod to rise so that the rat could move to the front and reach for the pellet. This was repeated until the rat began to quickly move to the front of the chamber to reach for the pellet after breaking the IR beam. At this point, the experimenter would stop baiting the rat to the rear of the chamber. Pre-training was complete once the rat requested a pellet and then immediately moved to the front to reach for the pellet 10 times.

#### Training

After pre-training, rats began 30-minute training sessions with the automated system. Rats were trained for 6 days per week until they reached stable performance (minimum of 35 reaches and a steady success rate above 40% over 3 sessions). Once behavioral criteria were met, rats were implanted with optical fibers.

### Optogenetics

Before testing with optogenetic interventions, rats were re-trained for 10 days while tethered to the patch cable without light delivery. This allowed rats to return to stable performance after surgery and adapt to the tether. During the 10 days of testing with optogenetic interventions, light was delivered on every trial at one of two different times. For “during reach” stimulation, the laser turned on when the rat broke the IR beam at the back of the chamber and remained on until 3 seconds after the video trigger event. For “between reach” stimulation, light was delivered beginning 5 seconds after the video trigger and remained on for 4 seconds (Figure 1D). The duration of “between reach” stimulation was approximately matched to the average duration of “during reach” stimulation. For ChR2- and EYFP-injected rats, 473 nm laser light (Opto Engine DPSS laser) was delivered at 20 Hz and an estimated 20 mW at the fiber tip based on pre-implantation measurements using a calibrated photodiode (Thorlabs S121C connected to Thorlabs PM100D Power Meter). The laser was on continuously, with 20 Hz stimulation achieved using an optical chopper (Thorlabs MC1F10HP) to eliminate transient power fluctuations as the laser is turned off and on. For Arch-injected rats, 532 nm laser light (Opto Engine DPSS laser) was delivered continuously at an estimated 20 mW at the fiber tip.

Following optogenetic testing, rats were tested for another 10 days with the patch cable attached to the implanted fiber and the laser activated. However, the patch cable-implanted fiber junction was physically occluded by inserting a piece of dense foam within the connector that holds the patch cable and optical fiber. Full occlusion of the laser was checked before each session by measuring light output at the fiber tip using a calibrated photodiode (Thorlabs S121C connected to Thorlabs PM100D Power Meter). In this way, all sensory cues were identical (e.g., visible light, optical shutter sounds) but light could not penetrate into the brain. The timing of light delivery was identical to that used during testing with optogenetic interventions.

### Analysis of Skilled Reaching Data

Analyses were performed using custom-written scripts and functions in MATLAB 2019a (MathWorks).

#### Number of trials and success rate

Reach outcome was scored by visual inspection as follows: 0 – no pellet presented or other mechanical failure; 1 – first trial success (obtained pellet on initial limb advance); 2 success (obtained pellet, but not on first attempt); 3 – forelimb advanced, pellet was grasped then dropped in the box; 4 – forelimb advance, but the pellet was knocked off the shelf; 5 – pellet was obtained using its tongue; 6 – the rat approached the slot but retreated without advancing its forelimb or the video triggered without a reach; 7 – the rat reached, but the pellet remained on the shelf; 8 – the rat used its contralateral paw to reach; 9 – laser fired at the wrong time; or 10 – used preferred paw after obtaining or moving pellet with tongue.

First reach success was calculated for each session by dividing the total number of scores of 1 by the total number of reaches (sum of scores of 1, 2, 3, 4, and 7). For both number of trials per session and first reach success rate, a baseline score was calculated for each rat by averaging the scores of the last two retraining sessions (Figure 2A,D). Number of trials and success rates for each session within “laser on” and “occlusion” sessions were normalized by dividing the score for that session by the averaged baseline score (Figures 2B-C, 2E-F, 3A-B, and 4A-D).

To assess how success rate changed within individual sessions, a moving average was calculated as the fraction of “1” scores in a moving block of 10 reaches. For averages within a group, the last data point for each individual was carried forward to the maximum number of reaches for any rat in that session. This avoided sudden changes in the average caused by dropout (Figures 2G and supplementary 1, 3C and supplementary 1, 4E and supplementary 1-2).

#### 3-dimensional reconstruction of reach trajectories

Bodyparts/objects identified in the direct and mirror views were triangulated to 3-dimensional points using custom MATLAB software (Bova et al., 2019). Prior to each session, several images of a cube with checkerboards (4 x 4 mm squares) on its sides were taken so that the checkerboards were visible in the direct and mirror views. These images were used to determine the essential matrix relating the direct and mirror views, which was used to determine how the real camera and “virtual” camera behind the mirror were translated and rotated with respect to each other (Hartley & Zisserman, 2003). By assuming a 3-dimensional coordinate system centered at the camera lens with the z-axis perpendicular to the lens surface, camera matrices were derived for the real and virtual cameras. These matrices were used to triangulate matching points in the camera and mirror views using the MATLAB triangulate function in the Computer Vision toolbox. 3-dimensional points with large reprojection errors were excluded from the analysis, which could happen if an object was identified accurately in one view but misidentified in the other.

#### Processing Reach Kinematics

To place reach kinematics in a common reference frame, the pellet location prior to reaching was identified and set as the origin. For left-pawed reaches, x-coordinates were negated to allow direct comparison with right-pawed reaches. The initial reach on each trial was identified by finding the first frame in which digits were visible outside the box, and then looking backwards in time until the paw started moving forward. The end of a reach was defined as the frame at which the tip of the second digit began to retract (“maximum reach extent”, z_digit2_). “Aperture” was calculated as the Euclidean distance between the tips of the 1^st^ and 4^th^ digits (in frames for which both were visible or could be estimated based on epipolar geometry). “Orientation” was calculated as the angle between a line connecting the 1^st^ and 4^th^ digits and a horizontal line (for left-pawed rats, orientation was calculated using the negated x-values to compare with right-pawed rats). Paw velocity was calculated as the Euclidean distance between the dorsum of the reaching paw in consecutive frames divided by the inter-frame interval (1/300 s).

#### Within-session kinematics

To assess how reach kinematics (i.e., maximum reach extent, aperture, paw orientation, and maximum reach velocity) changed within individual sessions, a moving average was calculated by averaging kinematic data across a moving block of 10 trials. For averages within an experimental group, the last data point was carried forward to the end of the data set. This avoided sudden changes in the average caused by rats performing different numbers of trials within a session.

#### Analysis of reach-to-grasp coordination

To monitor aperture and paw orientation as a function of the z-coordinate of the tip of the second digit (z_digit2_, Figures 8, 9 and all Figure 9 supplements), the first reach of each trial was isolated. The 3-dimensional trajectory of each digit tip for the initial reach was interpolated using piecewise cubic hermite polynomials (*pchip* in MATLAB) so that the 3-dimensional location of each digit was estimated for z_digit2_ = +20.0, +19.9, +19.8… −14.9, −15.0 mm from the pellet (positive numbers are as the paw approaches the pellet, negative numbers are past the pellet). This allows us to average aperture and orientation as a function of paw advancement (assessed by z_digit2_).

Two rats from the “ChR2 During” group were excluded from the averaged aperture and paw orientation as a function of z_digit2_ (Figure 9B, E). The majority of these rats’ reaches during “laser sessions” were so short that there were not enough trials with full trajectories to produce a meaningful average (see Figure 9 – figure supplements 1-2 for analysis with all 6 rats).

To compare the evolution of aperture and paw orientation between retraining, laser, and occlusion sessions, we compared digit aperture and paw orientation at specific z_digit2_ values. For all groups except “ChR2 During” and “ChR2 Between”, we evaluated aperture and orientation at z_digit2_ = 1 mm past the pellet (z_digit2_ = −1 mm). Because rats frequently did not reach past the pellet when dopamine neurons were activated “during reach”, we analyzed aperture and paw orientation at z_digit2_ = 7 mm before the pellet (z_digit2 =_ +7 mm) for this group.

### Abnormal Involuntary Movements (AIMs) Testing

Rats underwent AIMs testing twice – one day before the first day of retraining and one day after the last day of occlusion sessions. Rats were attached to the patch cable and placed into a clear plexiglass cylinder (diameter = 8.3 in). Two mirrors were placed behind the chamber so that the animal was visible in all positions in recordings. Once in the cylinder, animals underwent a series of 30 second stimulation epochs alternating with 30 second rest periods. Sessions always began with a rest period (baseline), and the order of laser power (estimated 5, 10, 15, 20, 25 mW at fiber tip) was randomly generated in Matlab. Stimulation was applied at 20 Hz at a 50% duty cycle. Stimulation sessions were video recorded at 60 frames-per-second (HC-V110, Panasonic).

AIMs videos were segmented into individual videos for each stimulation bout and assigned random codes so that scorers were blinded to the rat’s virus (ChR2 or EYFP), laser power, and day of testing. Axial and limb AIMs were scored for both severity (amplitude scale) and duration (basic scale) (Sebastianutto et al., 2016). The amplitude and basic scores were multiplied to create a composite score for axial and limb AIMs. Global AIMs scores were the sum of the axial and limb composite scores. Rotational behavior was also analyzed by counting the number of full 360 degree rotations in the contralateral and ipsilateral directions during each 30 second video. Ipsilateral turns were subtracted from contralateral turns to identify a rotational bias.

### Immunohistochemistry

Rats were deeply anesthetized with isoflurane (5%) and transcardially perfused with cold saline followed by 4% paraformaldehyde. Brains were post-fixed for no more than 24 h at 4°C, rinsed with saline, and moved through 20% and 30% sucrose solutions (in PBS) at 4°C. Sagittal sections (30 μm thickness) were taken around SNc and where the optical fiber was visible on a cryostat (Leica Microsystems). To verify localization of viral expression in dopamine neurons and optical fiber placement above SNc, we performed immunohistochemistry for tyrosine hydroxylase and EYFP. Mounted sections were washed with PBS and incubated with Triton X-100 and PBS (PBS-Tx) for 15 minutes. Slides were then incubated in 5% normal donkey serum (NDS) for 1 hour before primary antibody incubation (mouse anti-GFP, 1:1500, Life Technologies; rabbit anti-TH, 1:2000, Millipore) overnight at room temperature with NDS and PBS-Tx. Sections were then washed with PBS-Tx and incubated with secondary antibodies (Alexa Fluor 488 donkey anti-mouse, 1:500, Life Technologies; Alexa Fluor 555 donkey anti-rabbit, 1:500, Fisher Scientific) for 2 h at room temperature. After washing 4 times with PBS, sections were coverslipped with ProLong Diamond (Invitrogen), allowed to dry for 24 h, and then imaged with an Axioskops 2 Plus microscope fitted with an Olympus DP72 camera.

Images were stitched together and TH- and EYFP-stained images were overlaid in Photoshop to verify localization of viral expression to dopamine neurons. Images were evaluated by two people blinded to the behavioral outcomes of the individual rats on 1) sufficient virus expression in SNc and striatal dopamine neurons, and 2) location of fiber tip over SNc. Data from rats whose histology was evaluated as not meeting both of these criteria by both evaluators were removed from the analysis (n = 8 rats removed, Figure 1 – figure supplement 1). To obtain coordinates of optical fiber tips, histology images were overlaid on sagittal brain atlas images of the approximate M-L coordinate (Paxinos and Watson 1998) and A-P and D-V coordinates were ascertained.

### Statistics

Linear mixed-effects models were used to evaluate the effects of laser on performance outcomes and reach kinematics over sessions. We implemented linear mixed-effects models (using R *lmer*) with random intercepts/effects for each rat (where effect of laser varied between rats) and main interaction effects of group, session number, and laser. For normalized success rate and number of trials data, the inverse hyperbolic sine was taken before analysis in the linear mixed-effects model to deal with zeroes in the dataset. Post hoc contrast testing was performed on these linear mixed-effects models to make comparisons between specific sessions within groups (using R, ‘contest1D’). Similar models were used to evaluate changes in aperture and paw orientation at specific z_digit2_ coordinates in Figures 9C and 9F. However, random effects were designated where the effect of session varied between rats. To assess the effect of laser on reach kinematics in alternating sessions (Figure 10), we implemented a linear mixed-effects model with random intercepts/effects for each rat (where the effect of trial number within block varied between rats) and main interaction effects of laser and trial number within blocks. To assess how AIMs changed from the first to second day of testing and under different laser powers, we implemented a linear mixed-effects model with random effects for each rat (where the effect of test number varied between rats) and main interaction effects of group, test number, and laser power. To assess differences between groups in within-session analyses, we applied Wilcoxon rank sum tests (using MATLAB *ranksum*) at each trial number, with a *P* cutoff of 0.01 for significance.

## ACKNOWLEDGMENTS

This work was supported by the University of Michigan, an nVidia GPU grant, and NIH K08-NS072183.

## Video Captions

**Video 1** – Sample reach during the last “retraining” session of a “ChR2-During” rat showing the direct camera view, the mirror view of the paw dorsum, and 3D skeleton reconstruction. 2 trailing points are shown for each body part/object. Video is slowed 10x.

**Video 2** – Sample reach during the seventh “laser on” session for the same rat as in Video 1 showing the direct camera view, the mirror view of the paw dorsum, and 3D skeleton reconstruction. 2 trailing points are shown for each body part/object. Video is slowed 10x.

**Video 3** – Sample reach during the last “retraining” session of an “Arch-During” rat showing the direct camera view, the mirror view of the paw dorsum, and 3D skeleton reconstruction. 2 trailing points are shown for each body part/object. Video is slowed 10x.

**Video 4** – Sample reach during the tenth “laser on” session for the same rat as in Video 3 showing the direct camera view, the mirror view of the paw dorsum, and 3D skeleton reconstruction. While the reaches in Videos 3 and 4 are superficially similar, the rat reaches further past the pellet after repeated dopamine neuron inhibition. 2 trailing points are shown for each body part/object. Video is slowed 10x.

**Video 5 –** Sample reaches from a rat that received “during reach” stimulation in alternating trial blocks demonstrating that kinematic changes induced by dopamine neuron stimulation are enduring. Reach 1 – at baseline, the rat extends its paw past the pellet to grasp it. Reach 2 – after several reaches with stimulation, the second digit extends just to the pellet, which is knocked off the pedestal. Reach 3 – after more reaches with stimulation, the reach comes far short of the pellet. Reach 4 – with stimulation off, reach kinematics return to baseline. Reach 5 – on the next reach, stimulation is reinstated and kinematics are markedly abnormal.

**Video 6 –** Context-dependent AIMs. ChR2-injected rat showing AIMs (axial and limb dyskinesias) with dopamine neuron stimulation during the second day of AIMs testing. The same rat does not show AIMs when receiving the same stimulation parameters (estimated 20 mW at the fiber tip, 20 Hz) during a reach.

---

**Figure 1 – figure supplement 1.**
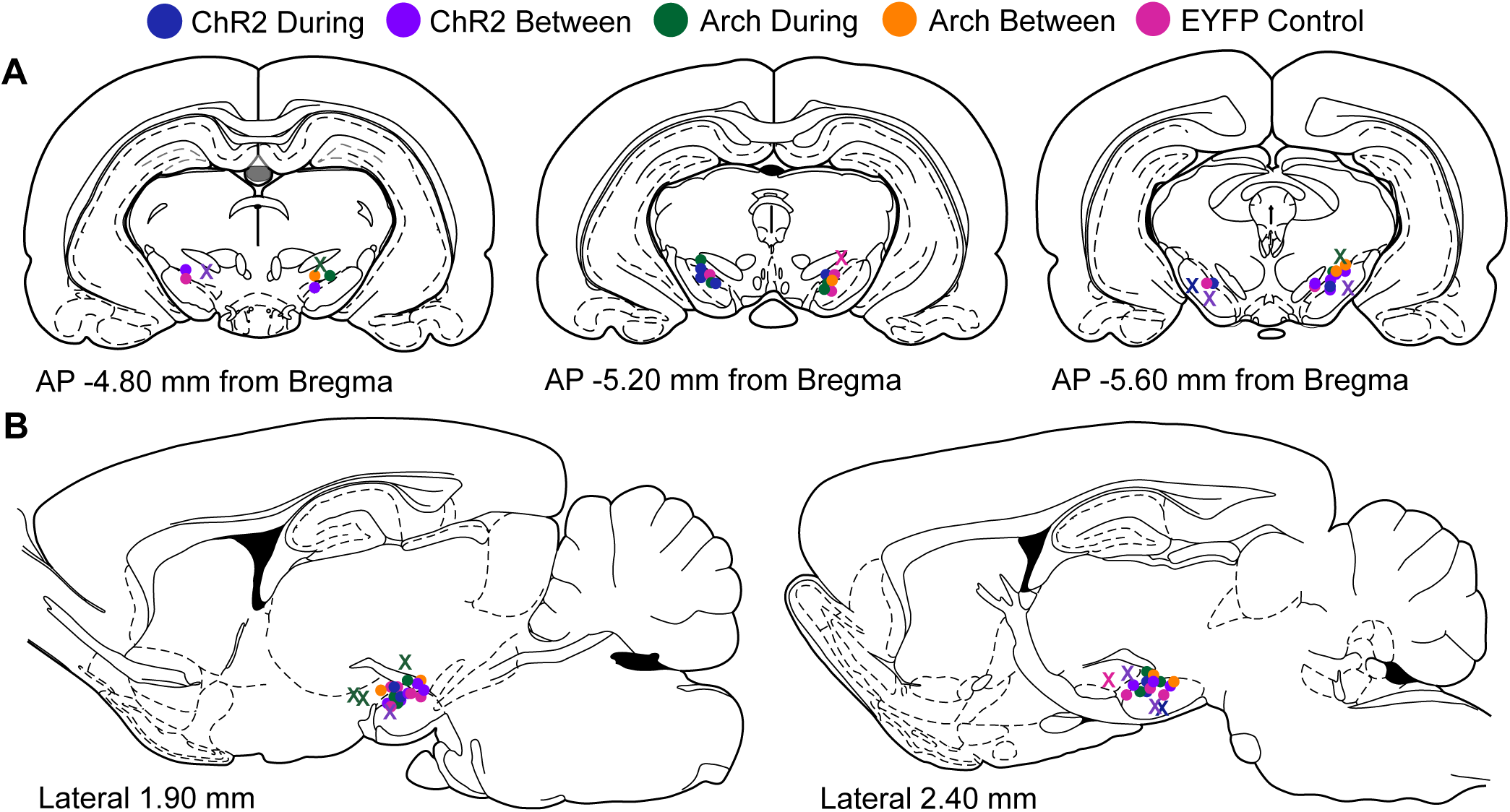
Optical fiber locations. **(A)** Circles indicate optical fiber tip locations in rats analyzed from each group superimposed on coronal rat brain atlas images (Paxinos and Watson 1998). X’s indicate optical fiber tip locations of rats excluded from the analysis due to either fiber misplacement or lack of opsin expression (see Materials and Methods). **(B)** Optical fiber tip locations superimposed on sagittal rat brain atlas images.

**Figure 2 – figure supplement 1.**
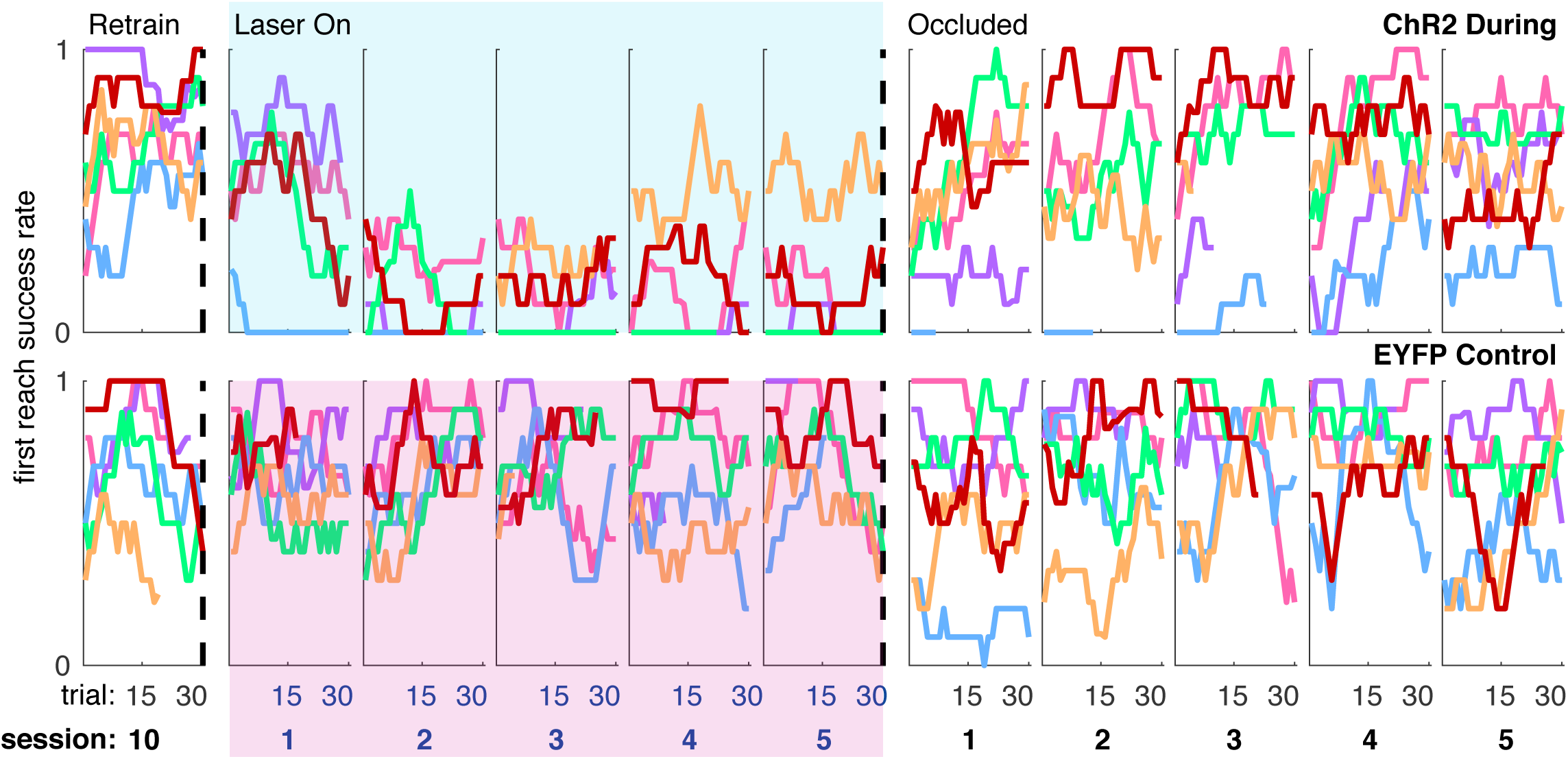
Moving average of success rate within individual sessions (ChR2 During and EYFP) for each rat. Each colored line represents the same rat across panels.

**Figure 3 – figure supplement 1.**
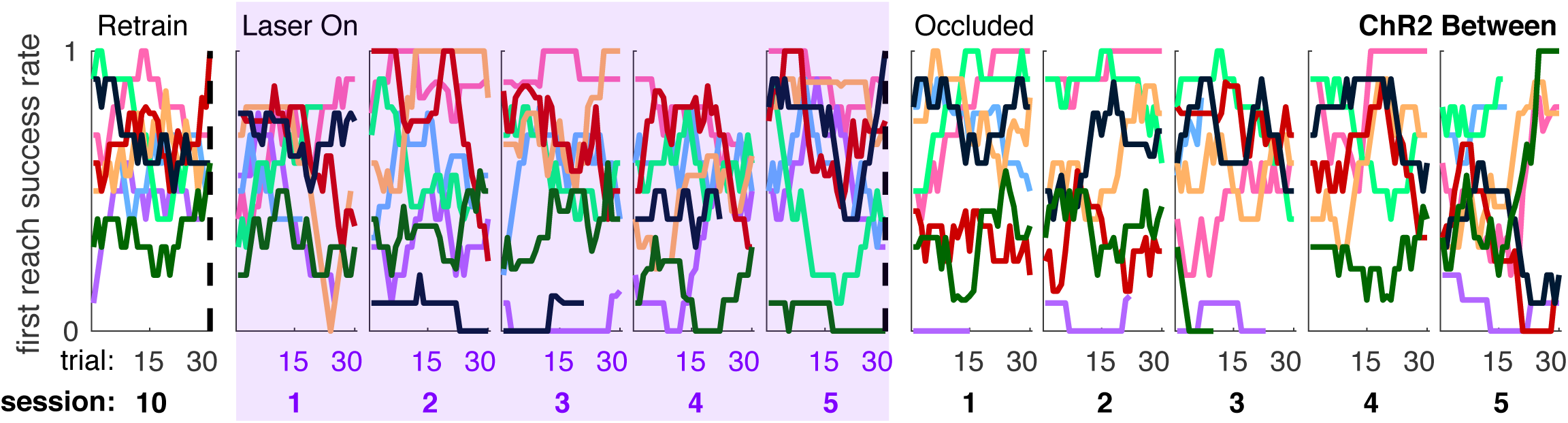
Individual rat data for moving average of success rate across trials within individual sessions (“ChR2 Between”). Each colored line represents the same rat across panels.

**Figure 4 – figure supplement 1.**
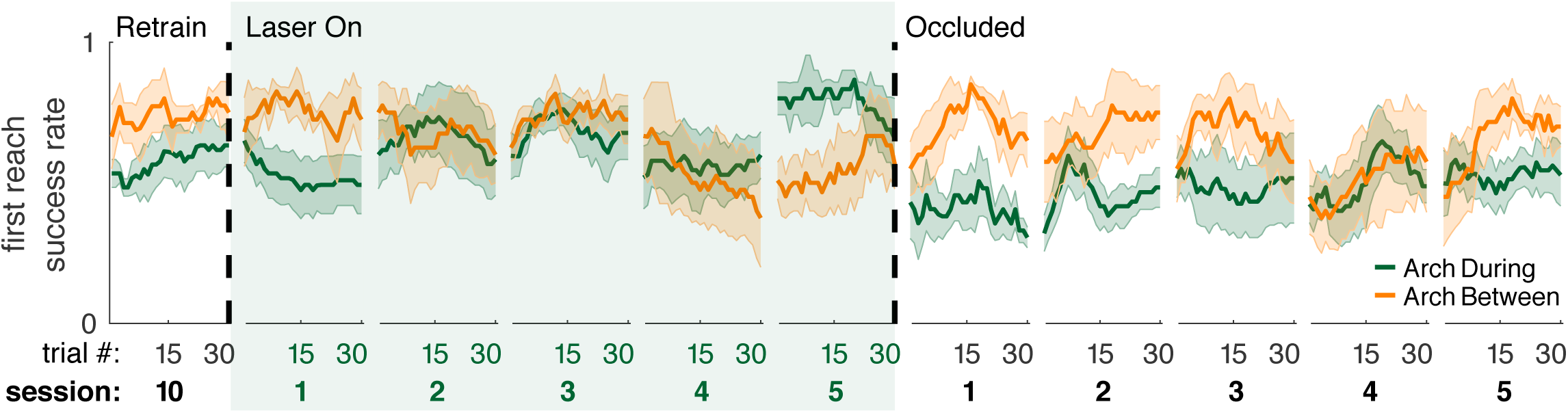
Moving average of success rate across trials within individual sessions in Arch-injected rats receiving dopamine neuron inhibition between reaches. “During reach” data from Figure 4 are shown for comparison.

**Figure 4 – figure supplement 2.**
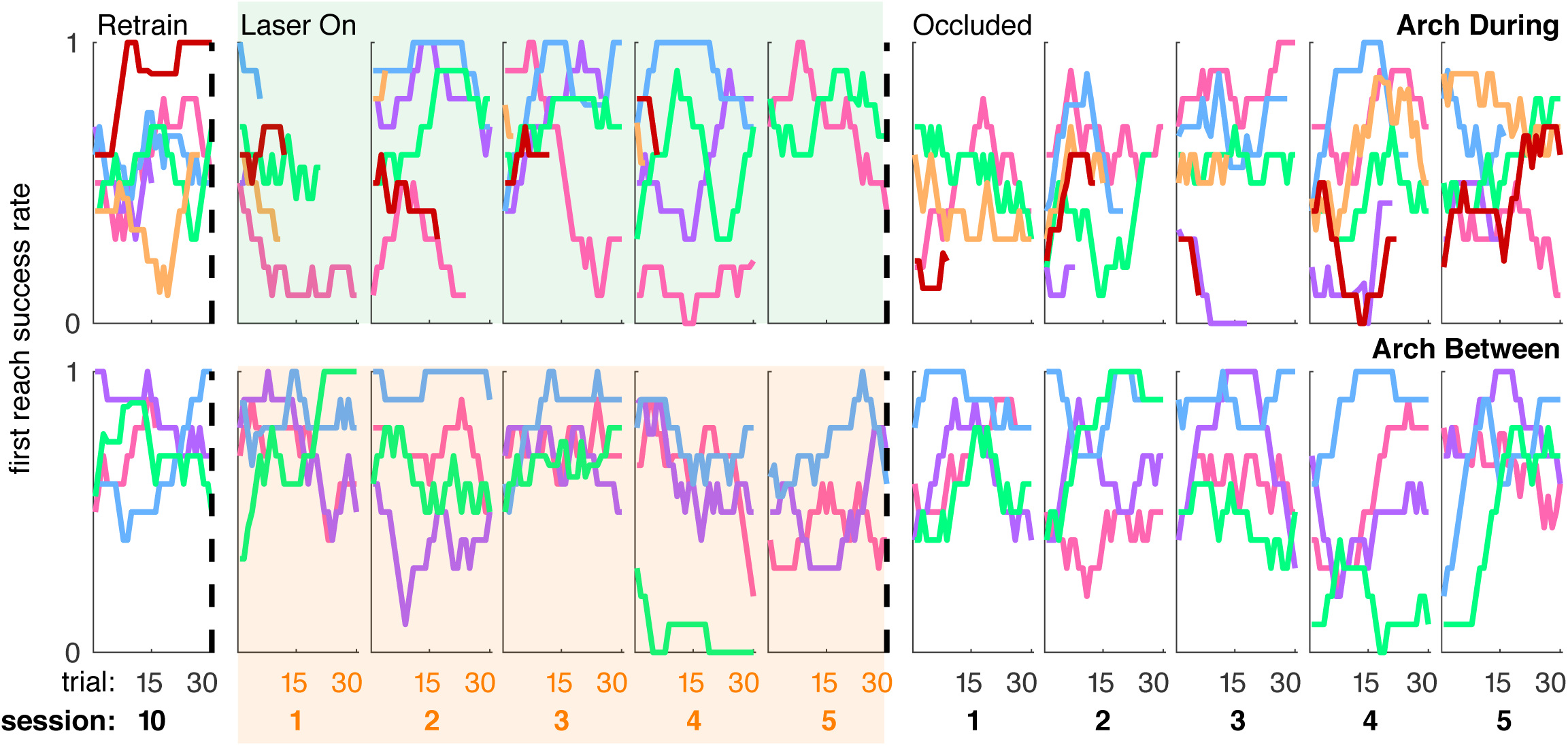
Individual rat data for moving average of success rate across trials within individual sessions (“Arch During” and “Arch Between”). Each colored line represents the same rat across panels.

**Figure 6 – figure supplement 1.**
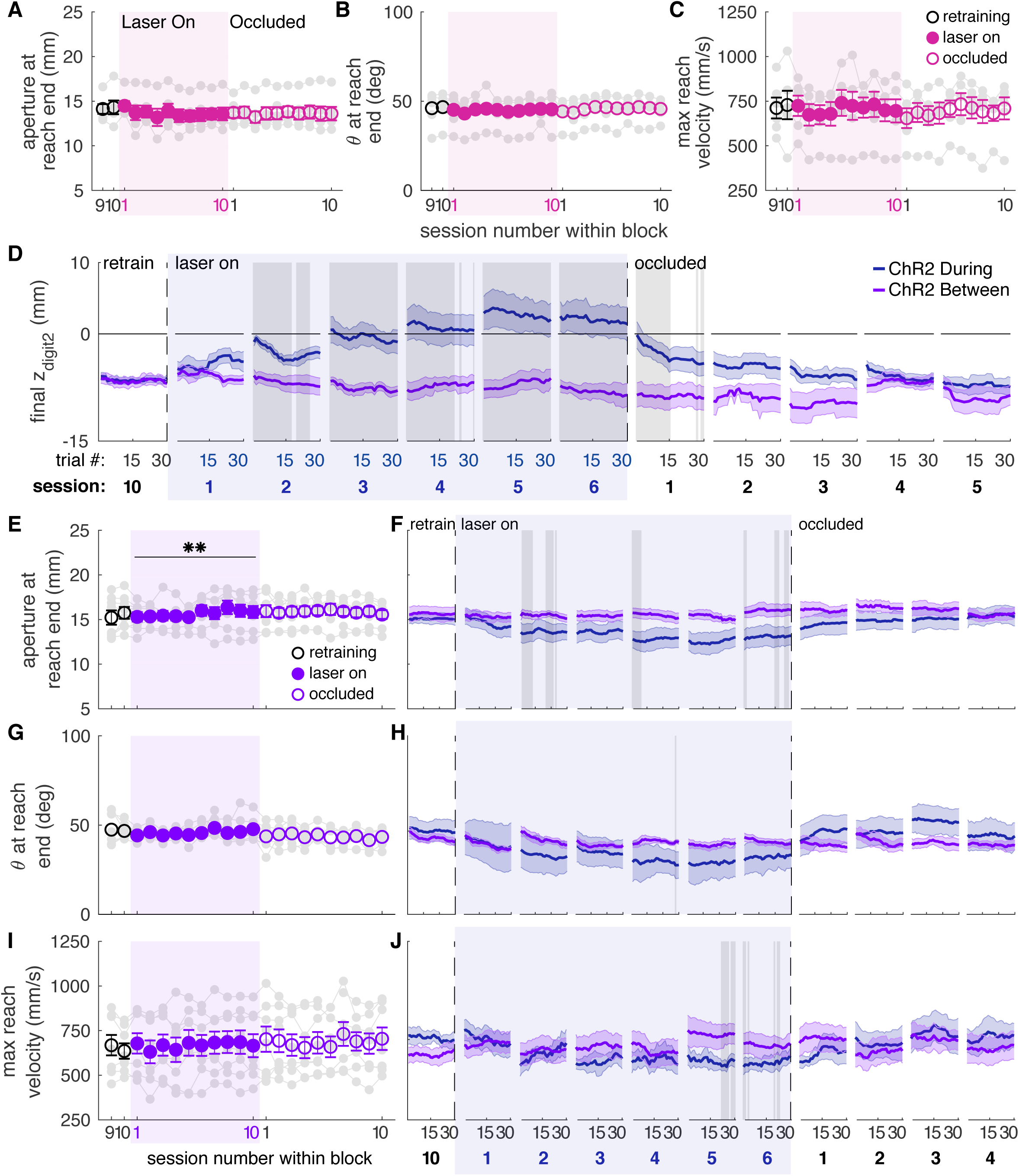
Reach-to-grasp kinematics do not change in EYFP control rats or ChR2-injected rats receiving between reach stimulation. **(A)** Average grasp aperture in EYFP control rats. Grey lines represent individual rats. Linear mixed model: effect of laser: *t*(48) = −0.16, *P* = 0.88; interaction between laser and session: *t*(585) = −1.14, *P* = 0.26. **(B)** Same as (A) but for paw orientation. Linear mixed model: effect of laser: *t*(75) = −0.65, *P* = 0.52; interaction between laser and session: *t*(585) = 0.47, *P* = 0.64. **(C)** Same as (A) and (B) but for maximum reach velocity. Linear mixed model: effect of laser: *t*(49) = −0.23, *P* = 0.82; interaction between laser and session: *t*(585) = 0.72, *P* = 0.47. **(D)** Moving average of maximum reach extent for “between reach” stimulation within the last “retraining” session, first 6 “laser on” sessions, and first 5 “occlusion” sessions. Grey shaded areas represent trials with a statistically significant difference between groups (Wilcoxon rank sum test, *P* < 0.01). Data for “during reach” stimulation from Figure 6D are shown for comparison. **(E)** Average grasp aperture for “between reach” stimulation. Linear mixed model: effect of laser: *t*(48) = −0.60, *P* = 0.55; interaction between laser and session: *t*(585) = 2.59, *P* = 9.76×10^-3^. **(F)** Moving average of aperture at grasp end within the last “retraining” session, first 6 “laser on” sessions, and first 4 “occlusion” sessions. Grey shaded areas represent trials with a statistically significant difference between groups (Wilcoxon rank sum test, *P* < 0.01). **(G)** Same as (E) for paw orientation. Linear mixed model: effect of laser: *t*(74) = −0.67, *P* = 0.51; interaction between laser and session: *t*(585) = 1.85, *P* = 0.06. **(H)** Moving average of paw angle at grasp end within the last “retraining” session, first 6 “laser on” sessions, and first 4 “occlusion” sessions. Grey shaded areas represent trials with a statistically significant difference between groups (Wilcoxon rank sum test, *P* < 0.01). **(I)** Same as (E) and (G) for maximum reach velocity. Linear mixed model: effect of laser: *t*(49) = 0.17, *P* = 0.87; interaction between laser and session: *t*(585) = −0.43, *P* = 0.67. **(J)** Moving average of maximum reach velocity within the last “retraining” session, first 6 “laser on” sessions, and first 4 “occlusion” sessions. Shaded colored areas in D, F, H, J and error bars in A, B, C, E, G, I represent s.e.m. ** indicates p < 0.01 for the laser-session interaction term in panel E.

**Figure 6 – figure supplement 2.**
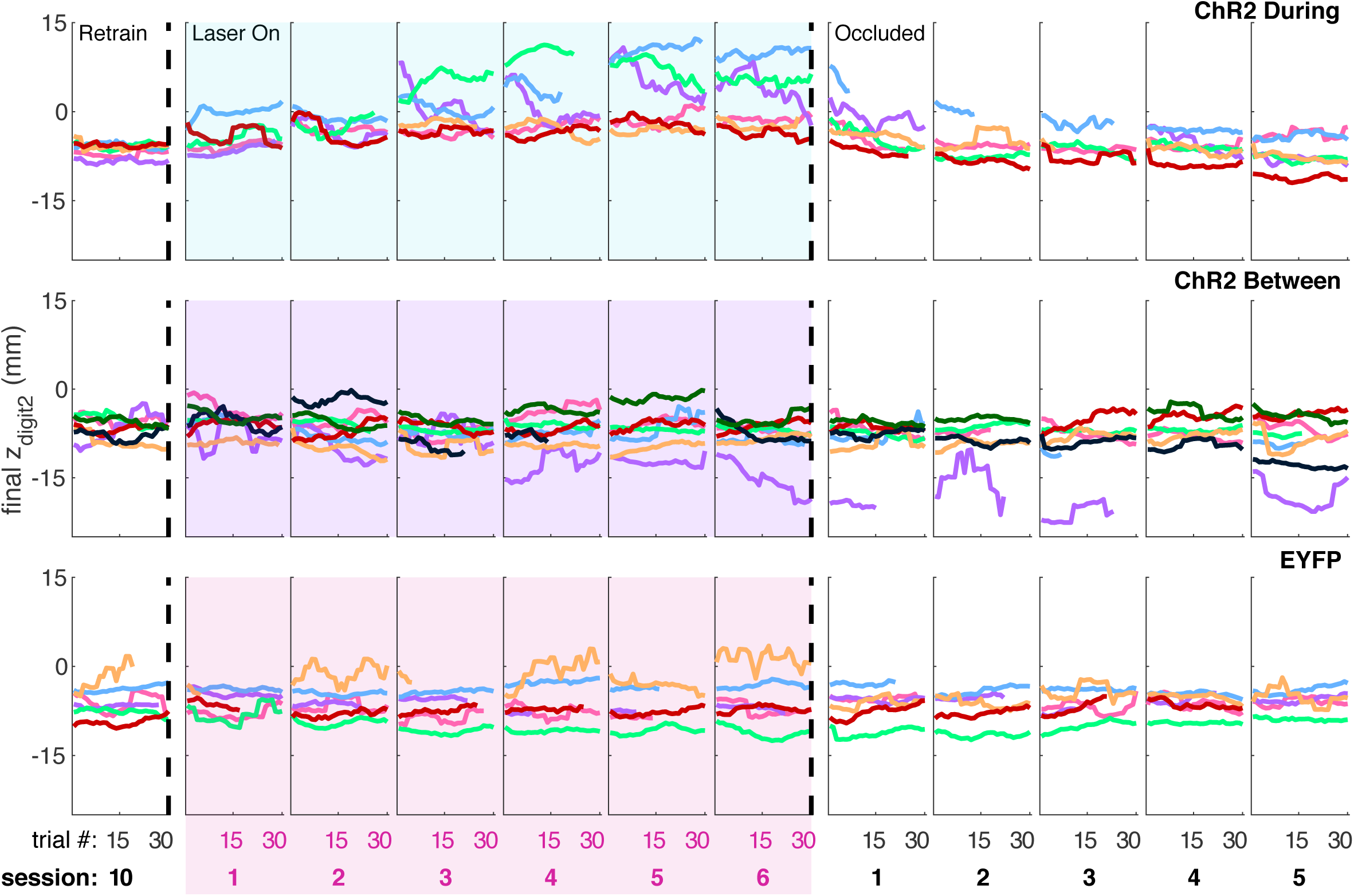
Individual rat data for moving average of maximum reach extent across trials within individual sessions (“ChR2 During”, “ChR2 Between” and “EYFP”). Each colored line represents the same rat across panels.

**Figure 6 – figure supplement 3.**
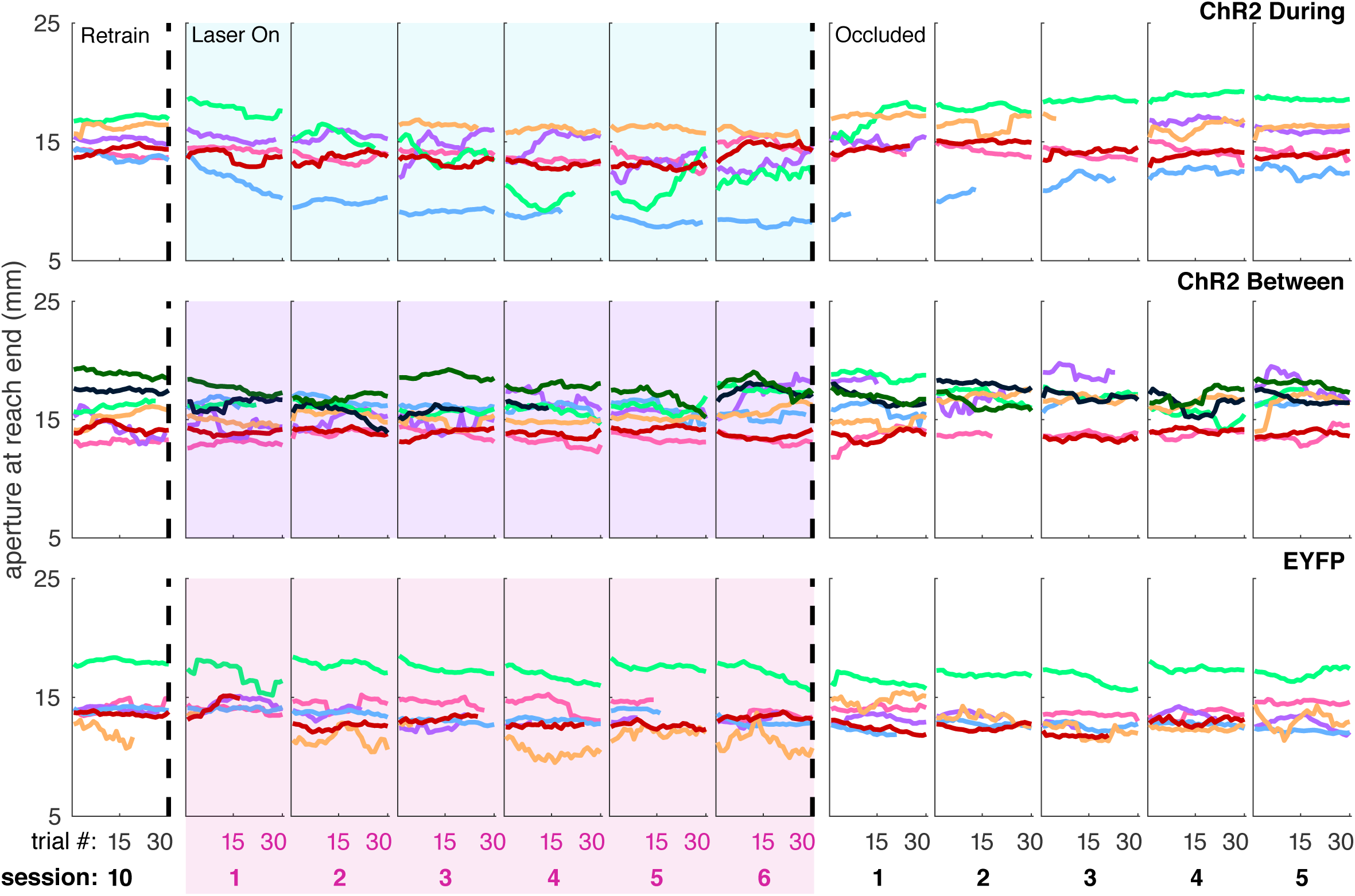
Individual rat data for moving average of grasp aperture across trials within individual sessions (“ChR2 During”, “ChR2 Between” and “EYFP”). Each colored line represents the same rat across panels.

**Figure 6 – figure supplement 4.**
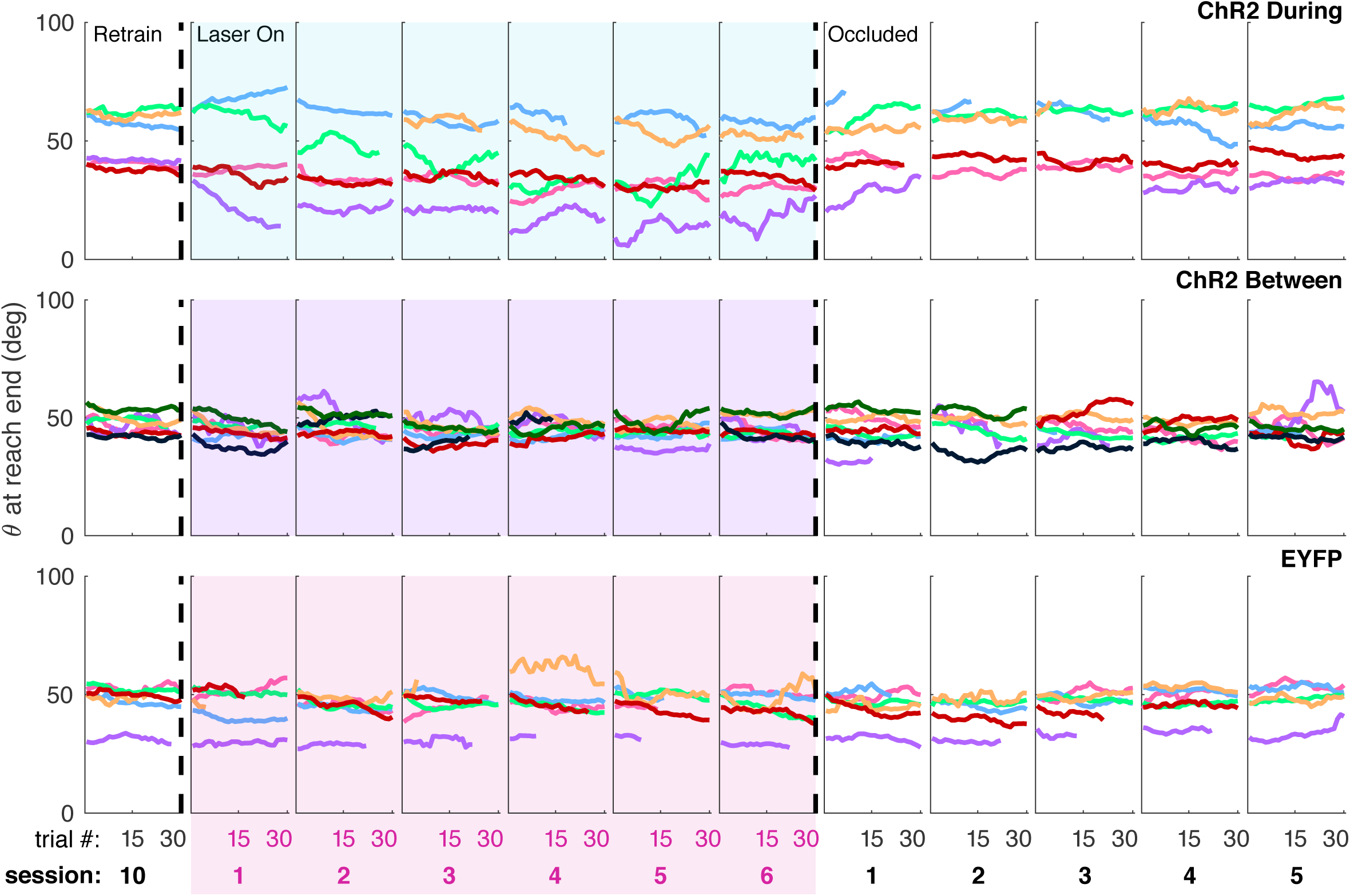
Individual rat data for moving average of paw angle across trials within individual sessions (“ChR2 During”, “ChR2 Between” and “EYFP”). Each colored line represents the same rat across panels.

**Figure 6 – figure supplement 5.**
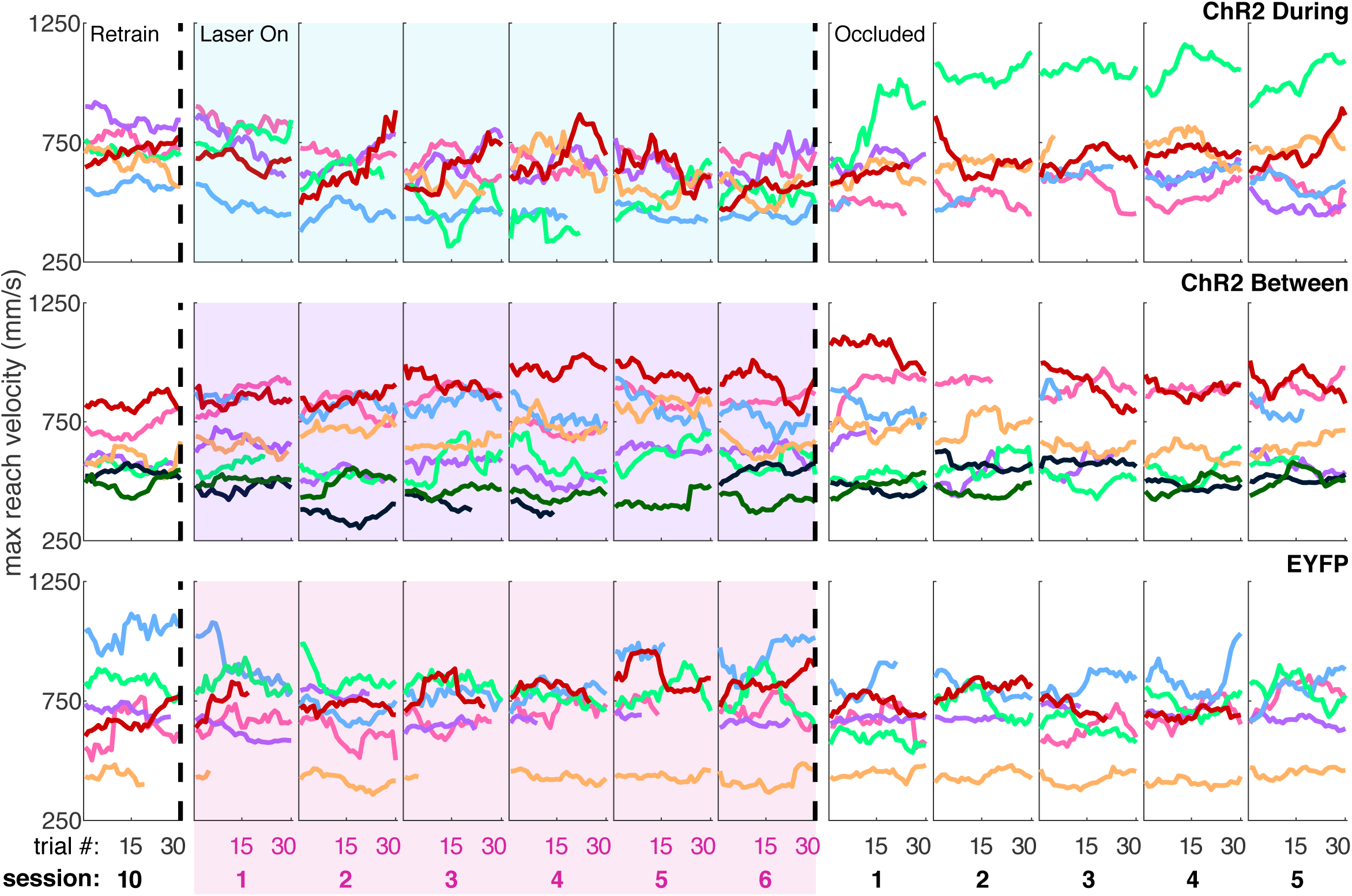
Individual rat data for moving average of maximum reach velocity across trials within individual sessions (“ChR2 During”, “ChR2 Between” and “EYFP”). Each colored line represents the same rat across panels.

**Figure 7 – figure supplement 1.**
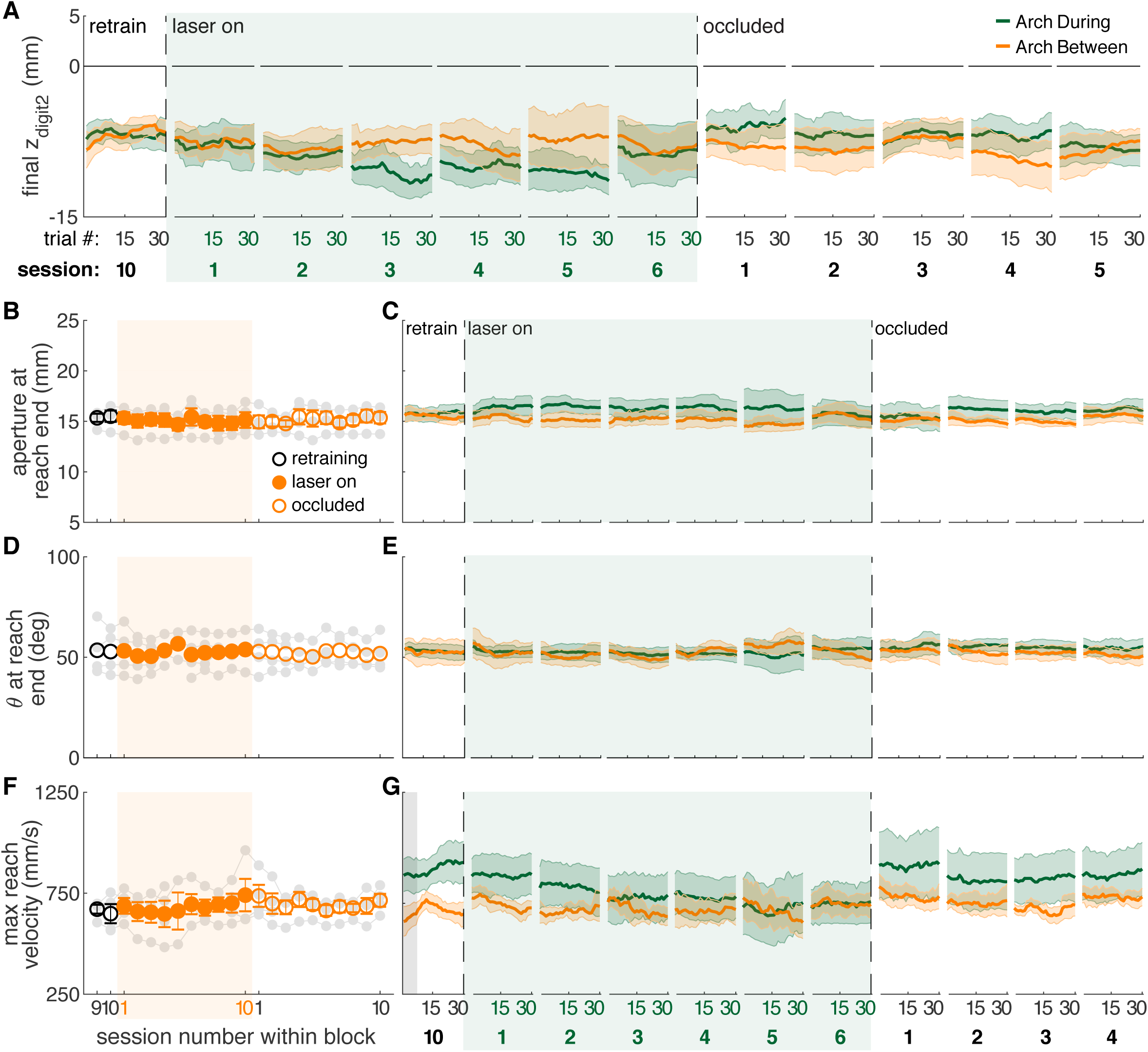
Reach-to-grasp kinematics do not change with dopamine neuron inhibition between reaches. **(A)** Moving average of maximum reach extent for “between reach” inhibition within the last “retraining” session, first 6 “laser on” sessions, and first 5 “occlusion” sessions. Data for “during reach” inhibition from Figure 7C are shown here for comparison. **(B)** Average aperture at reach end for “between reach” inhibition. Linear mixed model: effect of laser: *t*(48) = 0.05, *P* = 0.96; interaction between laser and session: *t*(585) = 0.26, *P* = 0.80. **(C)** Moving average of aperture at reach end for “between reach” inhibition within the last “retraining” session, first 6 “laser on” sessions, and first 4 “occlusion” sessions. **(D)** Same as (B) for paw orientation: effect of laser: *t*(75) = −0.11, *P* = 0.91; interaction between laser and session: *t*(585) = 0.50, *P* = 0.62. **(E)** Moving average of paw angle at reach end for “between reach” inhibition within the last “retraining” session, first 6 “laser on” sessions, and first 4 “occlusion” sessions. **(F)** Same as (B) and (D) for maximum reach velocity: effect of laser: *t*(49) = −0.19, *P* = 0.85; interaction between laser and session: *t*(585) = 0.49, *P* = 0.62. **(G)** Moving average of maximum reach velocity at reach end for “between reach” inhibition within the last “retraining” session, first 6 “laser on” sessions, and first 4 “occlusion” sessions. Grey shaded areas represent trials with a statistically significant difference between groups (Wilcoxon rank sum test, *P* < 0.01). Shaded colored areas in A, C, E, G and error bars in B, D, F represent s.e.m.

**Figure 7 – figure supplement 2.**
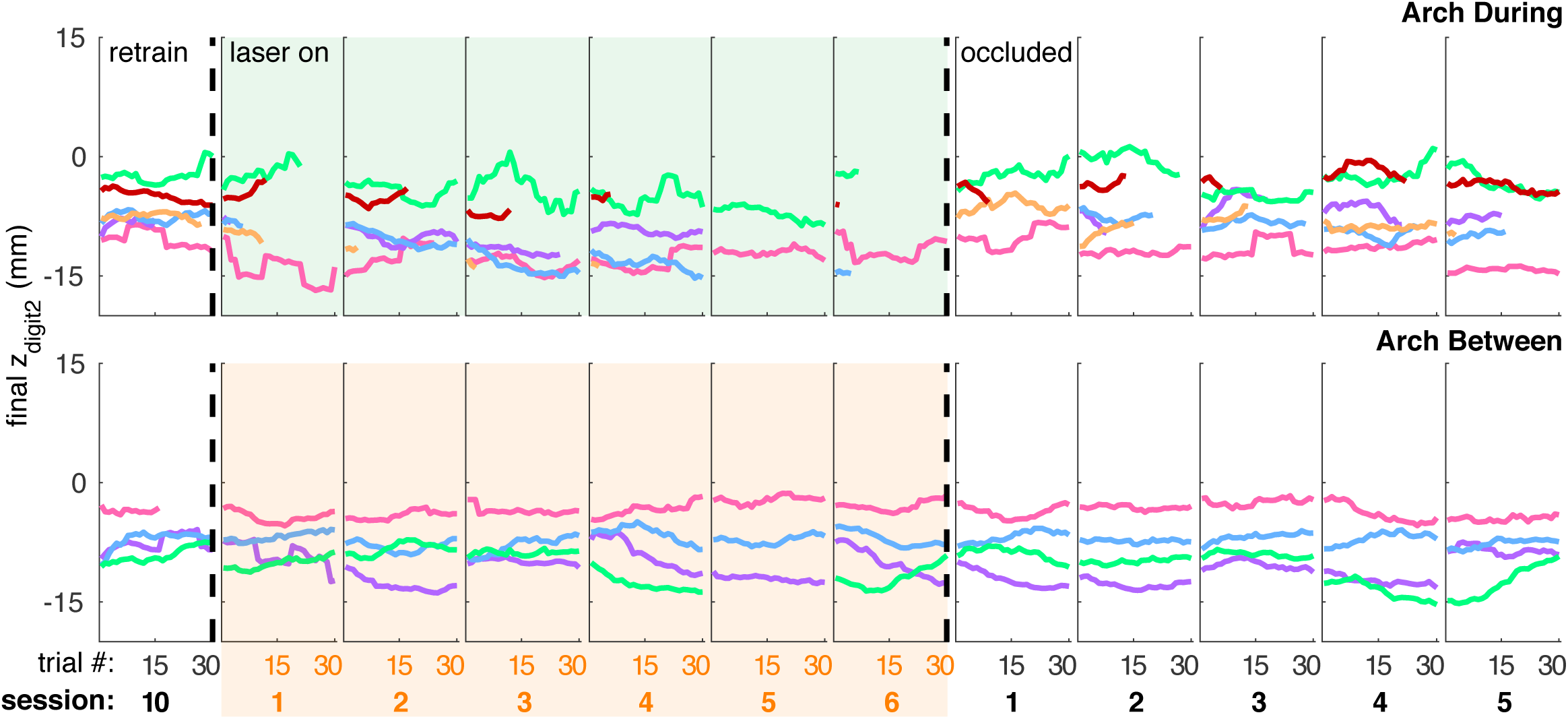
Individual rat data for moving average of maximum reach extent within individual sessions (“Arch During” and “Arch Between”). Each colored line represents the same rat across panels.

**Figure 7 – figure supplement 3.**
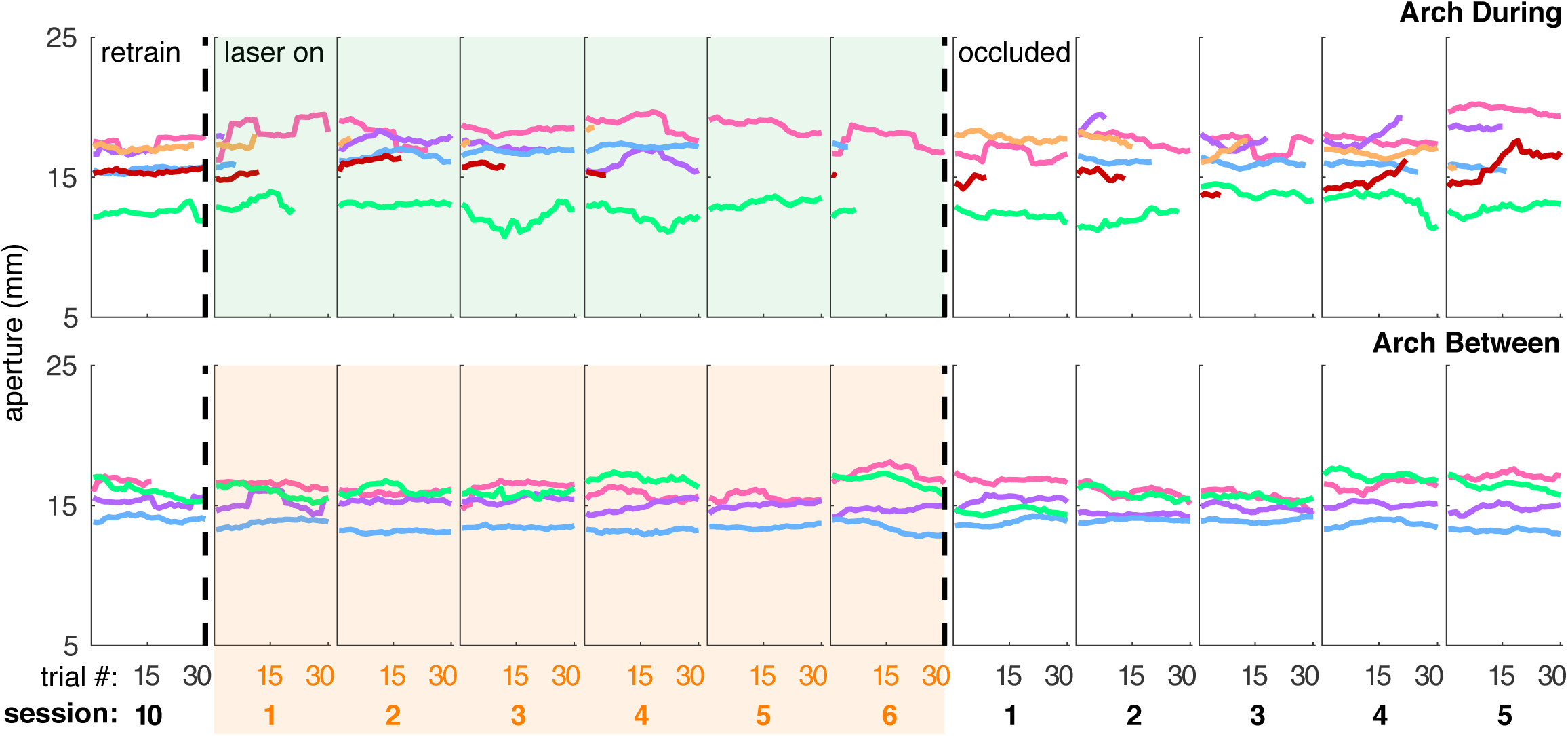
Individual rat data for moving average of grasp aperture at reach end within individual sessions (“Arch During” and “Arch Between”). Each colored line represents the same rat across panels.

**Figure 7 – figure supplement 4.**
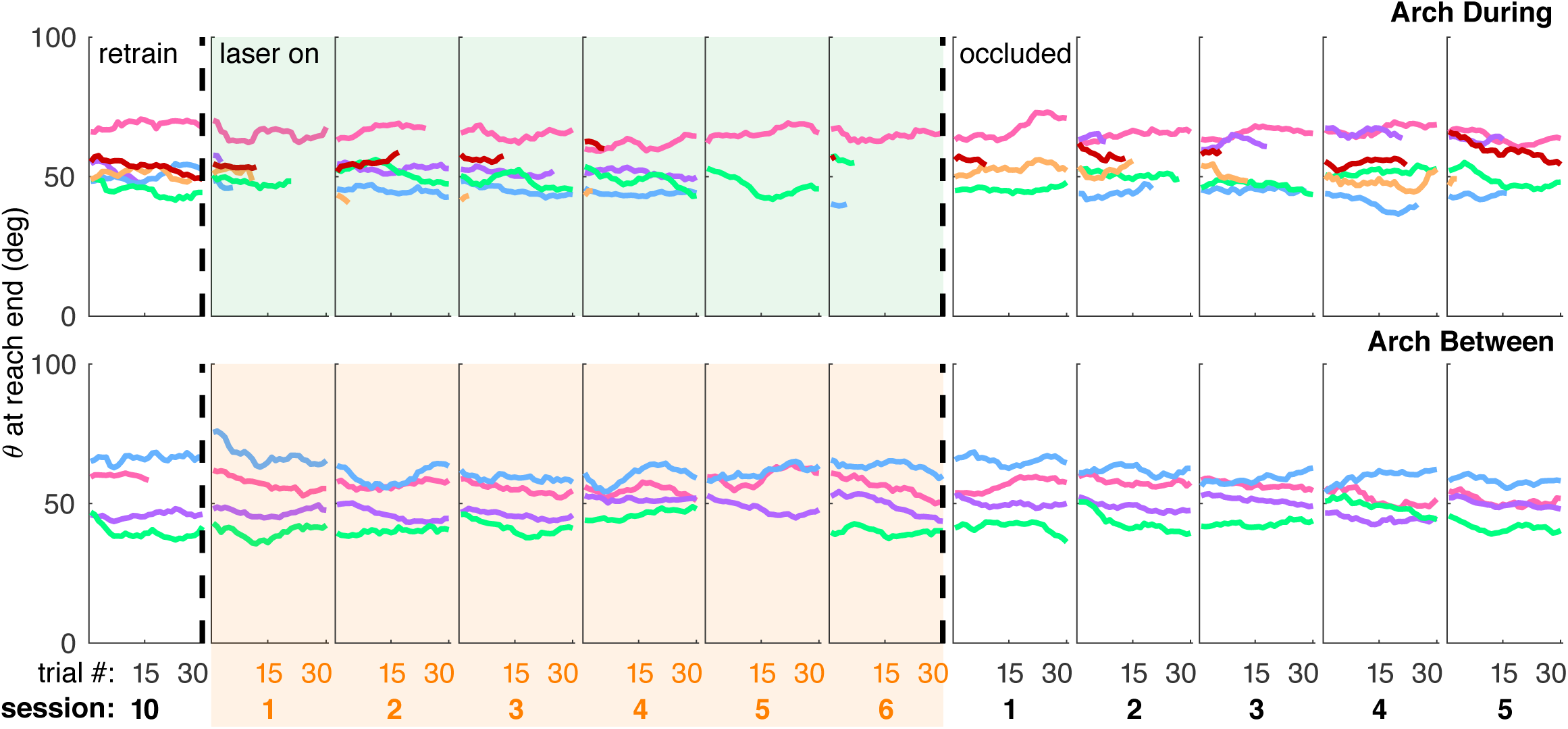
Individual rat data for moving average of paw angle at reach end within individual sessions (“Arch During” and “Arch Between”). Each colored line represents the same rat across panels.

**Figure 7 – figure supplement 5.**
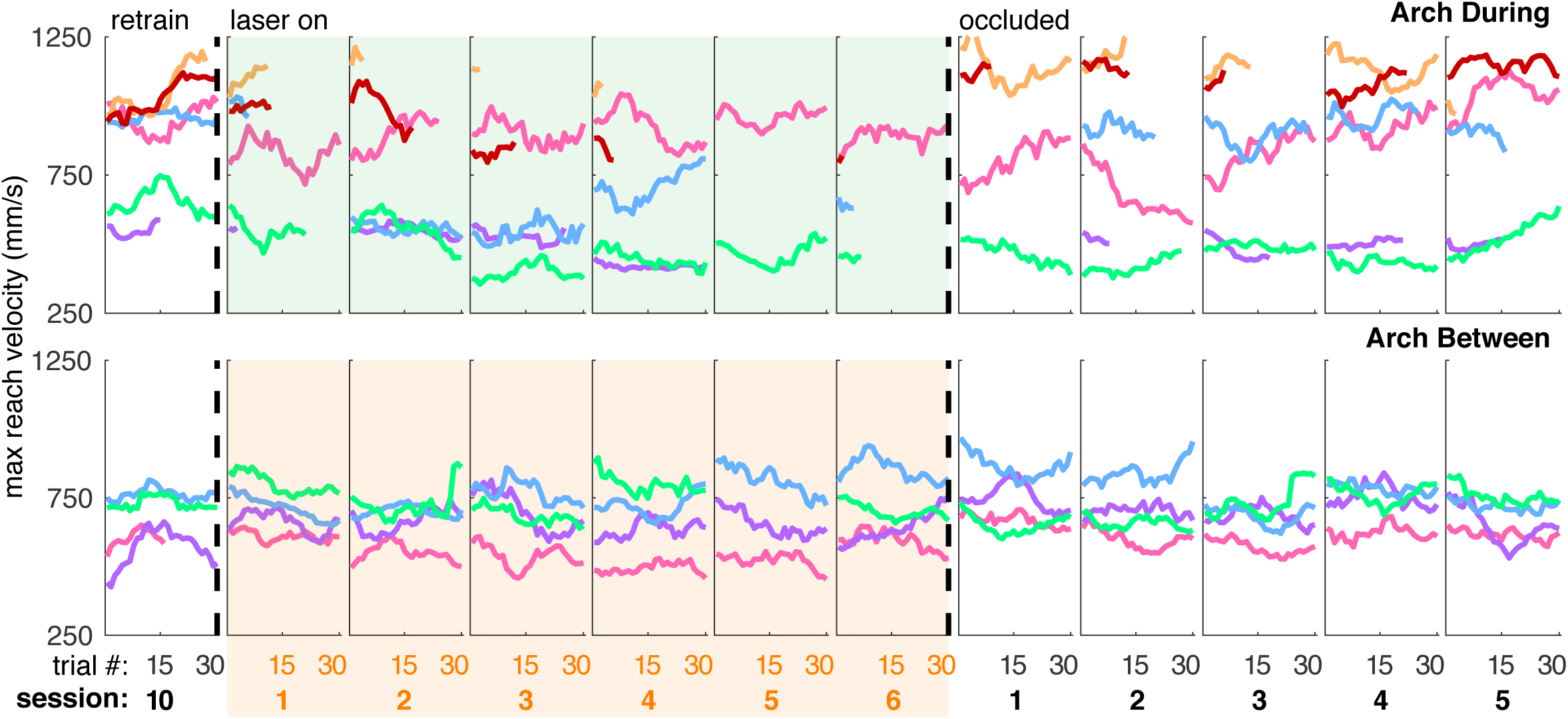
Individual rat data for moving average of maximum reach velocity within individual sessions (“Arch During” and “Arch Between”). Each colored line represents the same rat across panels.

**Figure 9 – figure supplement 1.**
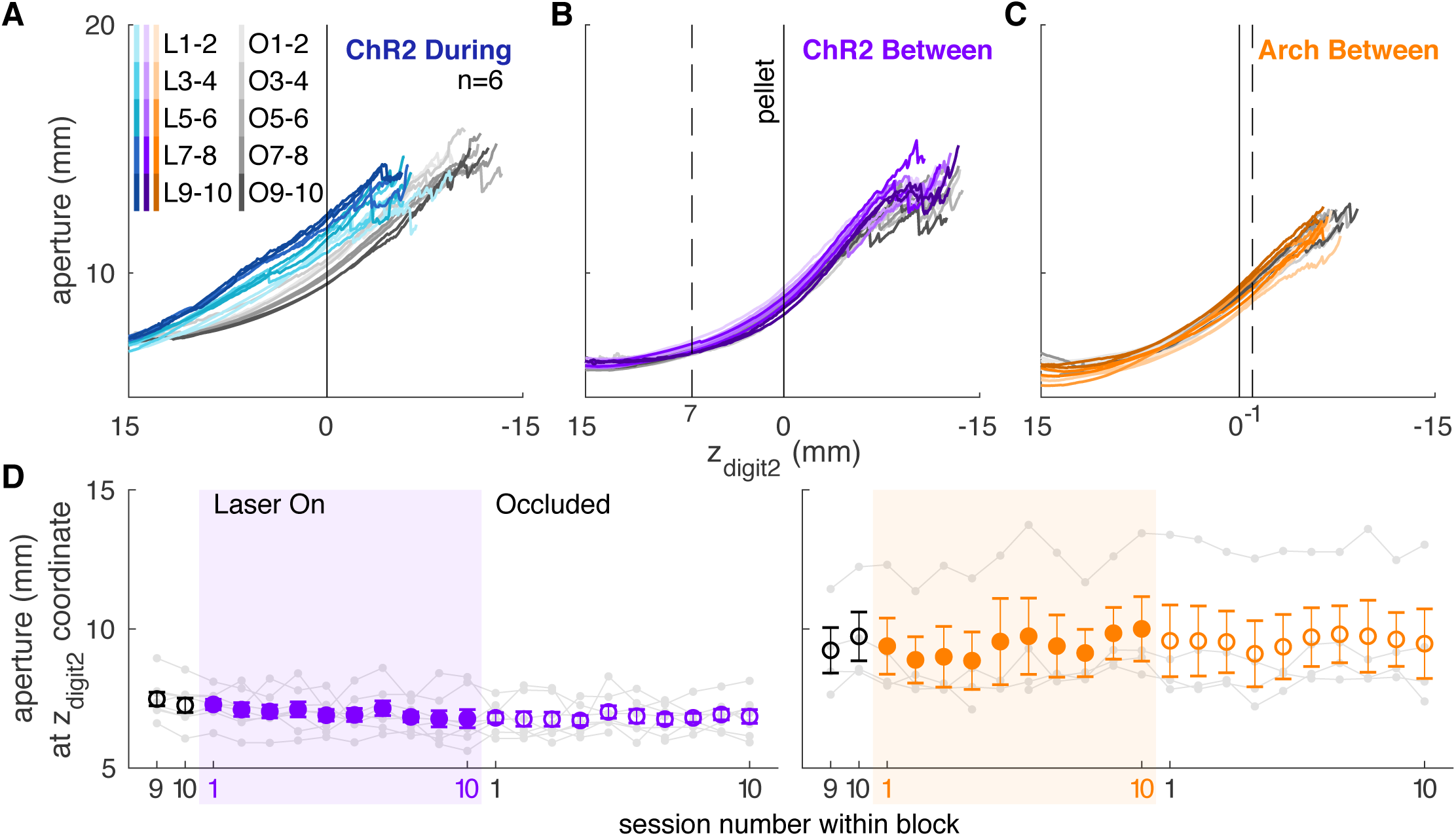
Dopamine manipulations between reaches do not affect reach-to-grasp coordination. **(A)** Mean aperture as a function of paw advancement in all “ChR2 During” rats (n = 6). L1-2, O1-2, … indicate laser on sessions 1-2, occlusion sessions 1-2, etc. **(B)** Mean aperture as a function of paw advancement for “between reach” stimulation. **(C)** Mean aperture as a function of paw advancement for “between reach” inhibition. **(D)** Average aperture at z_digit2_-coordinates indicated by dashed lines in (B) and (C) across all sessions. “Between reach” dopamine neuron stimulation did not affect aperture 7 mm from the pellet (linear mixed model: effect of laser: *t*(607) = 0.37, *P* = 0.71; interaction between laser and session: *t*(607) = 0.07, *P* = 0.94) or 1 mm past the pellet (linear mixed model: effect of laser: *t*(607) = 0.53, *P* = 0.60; interaction between laser and session: *t*(607) = −0.56, *P* = 0.58). Dopamine neuron inhibition between reaches did not affect aperture 1 mm past the pellet (linear mixed model: effect of laser: *t*(607) = −0.90, *P* = 0.37; interaction between laser and session: *t*(607) = 1.82, *P* = 0.07).

**Figure 9 – figure supplement 2.**
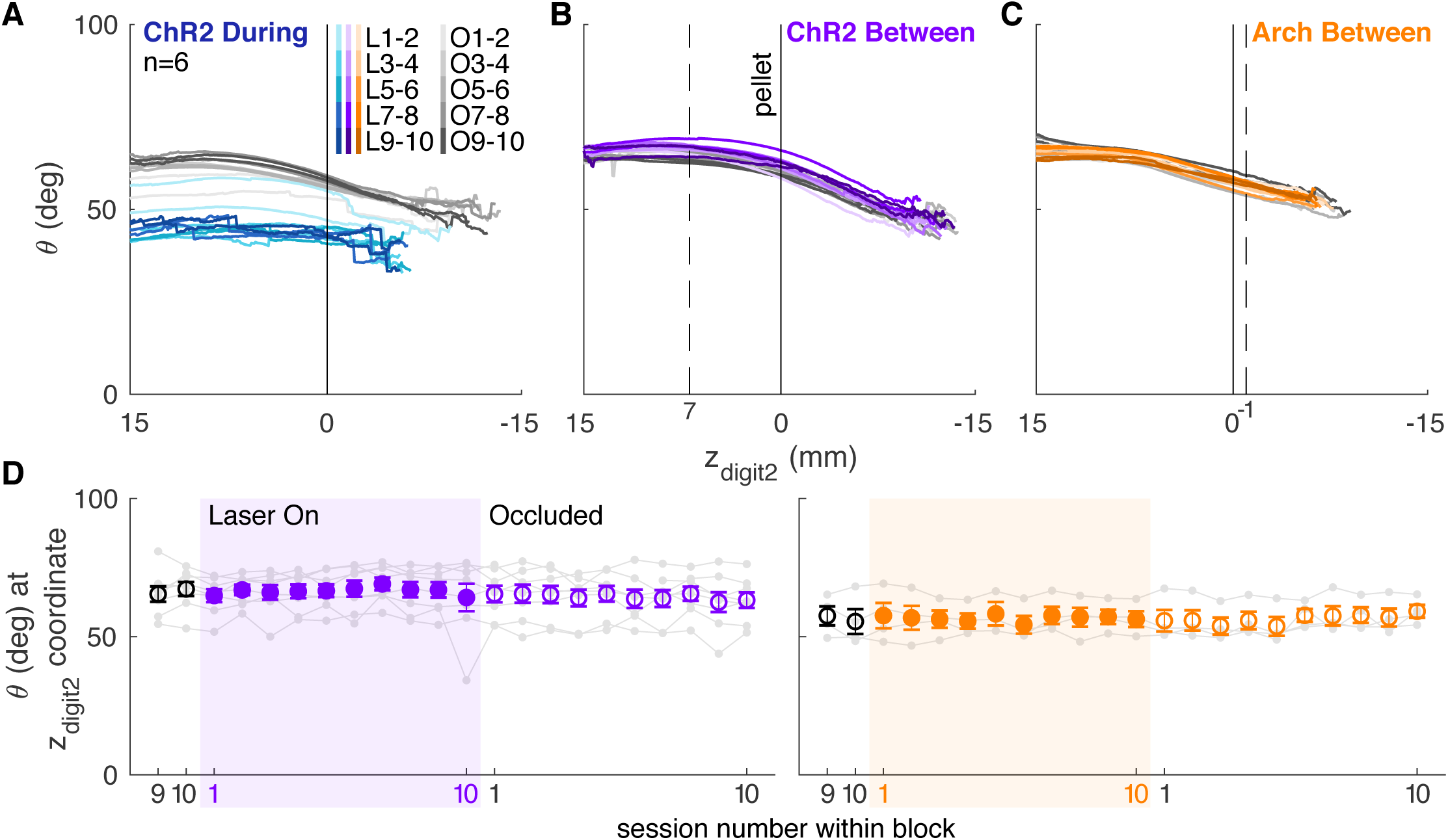
Dopamine manipulations between reaches do not affect coordination of reach-to-grasp kinematics (orientation). **(A)** Mean paw orientation as a function of paw advancement in all “ChR2 During” rats (n = 6). L1-2, O1-2, … indicate laser on sessions 1-2, occlusion sessions 1-2, etc. **(B)** Mean paw orientation as a function of paw advancement for “between reach” stimulation. **(C)** Mean paw orientation as a function of paw advancement for “between reach” inhibition. **(D)** Average paw orientation at z_digit2_-coordinates indicated by dashed lines in (B) and (C) across all sessions. “Between reach” dopamine neuron stimulation did not affect paw orientation at 7 mm away from the pellet (linear mixed model: effect of laser: *t*(607) = 0.27, *P* = 0.79; interaction between laser and session: *t*(607) = 0.22, *P* = 0.82) or 1 mm past the pellet (linear mixed model: effect of laser: *t*(607) = −0.41, *P* = 0.68; interaction between laser and session: *t*(607) = 1.17, *P* = 0.24). Dopamine neuron inhibition between reaches did not affect paw orientation at 1 mm past the pellet (linear mixed model: effect of laser: *t*(607) = 0.61, *P* = 0.54; interaction between laser and session: *t*(607) = −0.36, *P* = 0.72).

**Figure 9 – figure supplement 3.**
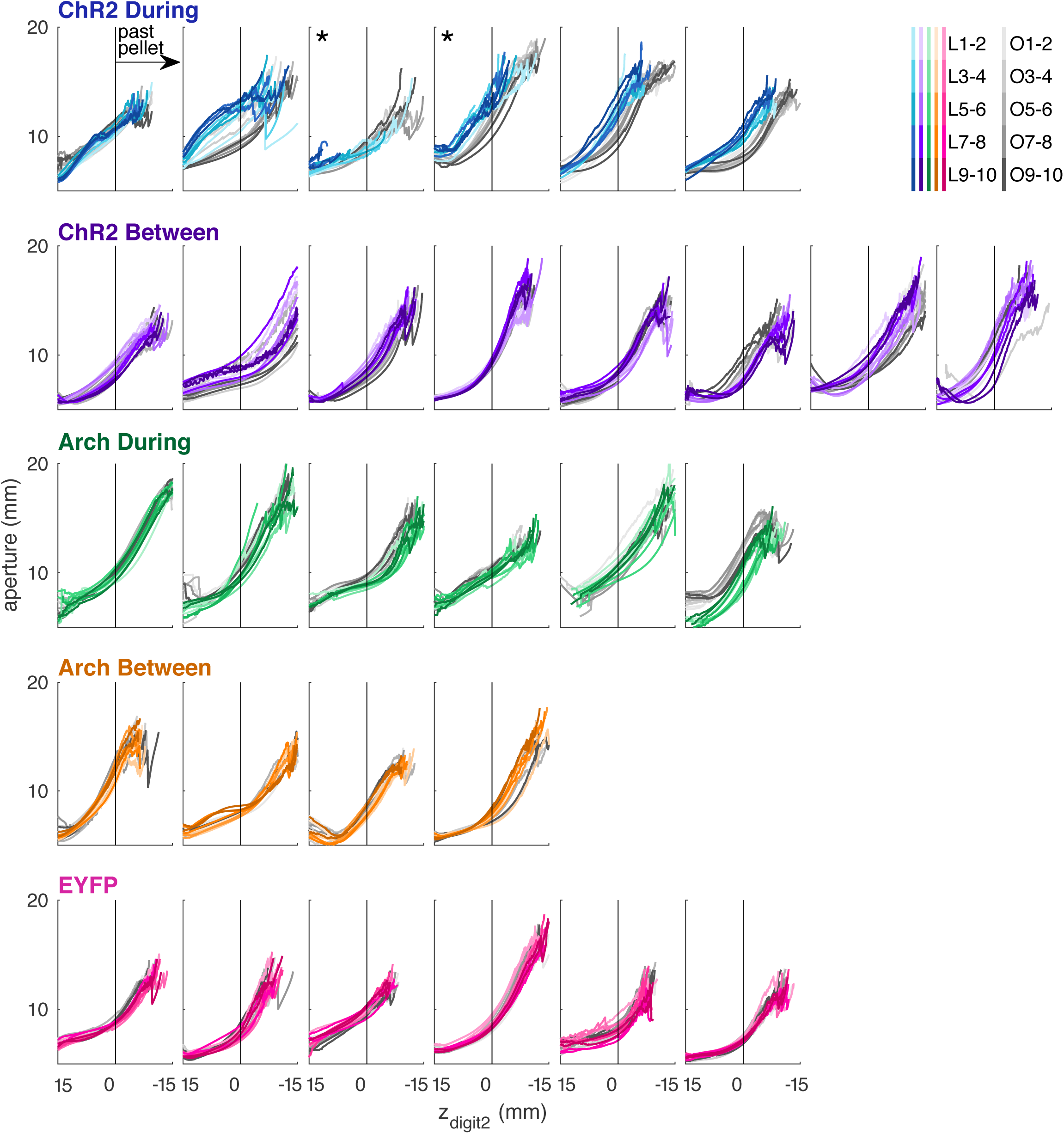
Mean aperture as a function of paw advancement for each rat. From top to bottom: ChR2 during reach stimulation, ChR2 between reach stimulation, Arch during reach inhibition, Arch between reach inhibition and EYFP during reach stimulation. * indicates rats that were excluded from averaged data in Figure 9B (see Methods). In the legend, L1-2, O1-2, … indicate laser on sessions 1-2, occlusion sessions 1-2, etc.

**Figure 9 – figure supplement 4.**
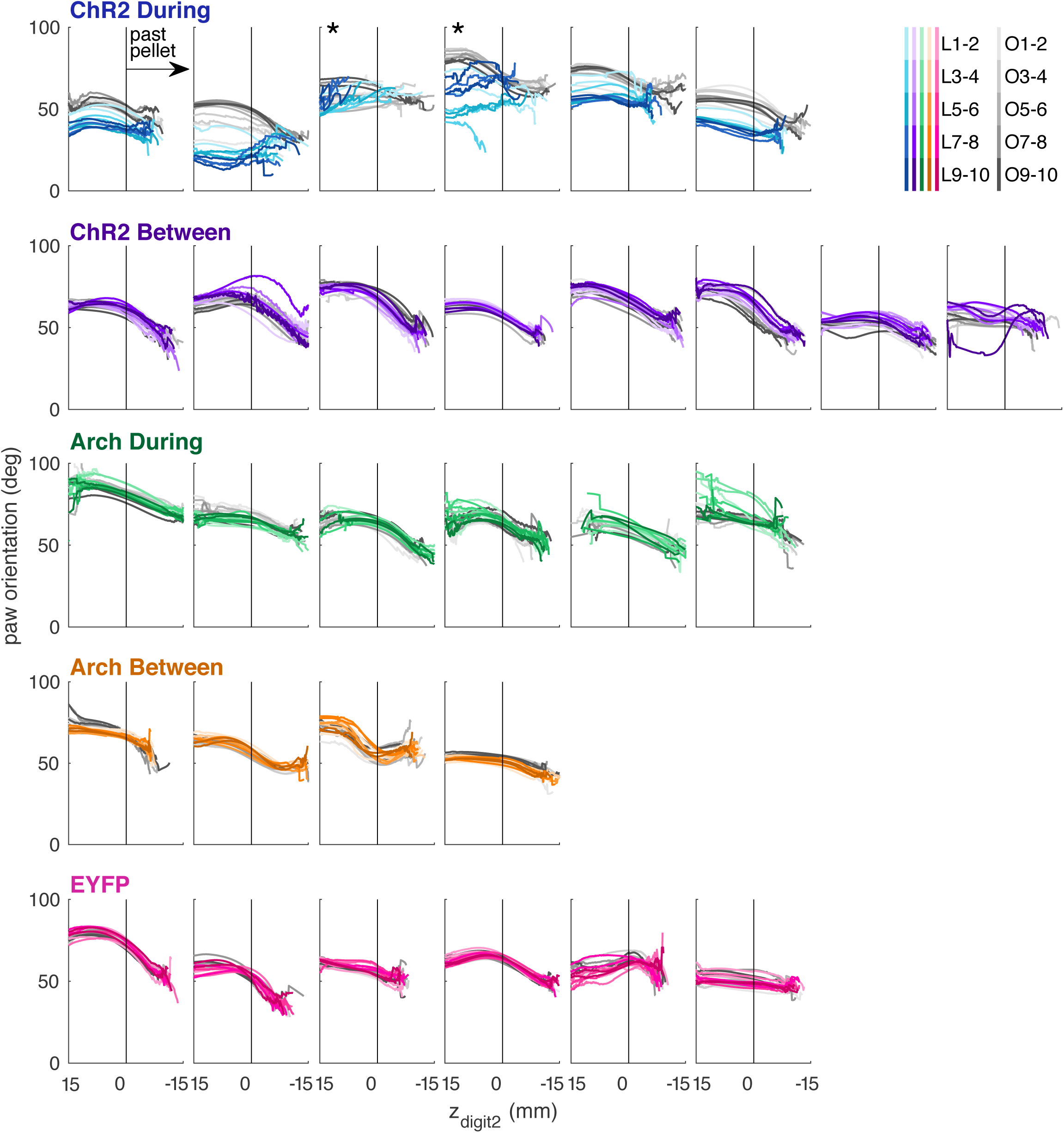
Mean paw orientation as a function of paw advancement for each rat. From top to bottom: ChR2 during reach stimulation, ChR2 between reach stimulation, Arch during reach inhibition, Arch between reach inhibition, and EYFP during reach stimulation. * indicates rats that were excluded from averaged data in Figure 9E (see Methods). In the legend, L1-2, O1-2, … indicate laser on sessions 1-2, occlusion sessions 1-2, etc.

